# Proteomics reveal temperature-coupled cobalamin homeostasis and pathogenicity in *Pseudomonas aeruginosa*

**DOI:** 10.64898/2026.01.27.701802

**Authors:** Viktoria Steck, Matthew R. McIlvin, Annie Stefanides, Eric A. Webb, Mak A. Saito

## Abstract

The opportunistic pathogen *Pseudomonas aeruginosa* is highly adaptable to different environmental conditions due to its versatile sensing and metabolic capabilities. Both external temperature and metal availability have a strong influence on the virulence and pathogenicity of *P. aeruginosa*, but the coupling between these two factors is not well understood. While iron is recognized as major player in nutritional immunity, the role of cobalt and the cobalt-containing vitamin B (cobalamin) during host infection remains unclear. Here, we investigate the environmental isolate *P. aeruginosa* PA254 using high-resolution global proteomics and cellular cobalamin measurements over a temperature gradient spanning environmental, host-associated, and heat-stress conditions (22–42 °C). PA254 occupies a continuum between an ambient-temperature state characterized by secreted virulence factors, exopolysaccharide-rich biofilms, and planktonic swimmers and surface swarmers; and a host-associated state characterized by a distinct set of virulence factors, alginate-dominated biofilms, and a strong proportion of surface twitching motility. Pathway analyses indicate a shift toward carbon sparing, energy conservation, redox control, and metabolic maintenance during a host-adapted lifestyle, along with the increased production of alternative iron acquisition strategies relying on heme and siderophores. Proteins of the cobalamin biosynthetic pathway declined significantly above ambient temperatures, despite constant intracellular B_12_ concentrations and sustained abundance of B_12_-dependent enzymes across all conditions. This decoupling of biosynthesis from cellular pools implies temperature-dependent changes in prioritization, biosynthetic efficiency, or recycling within B_12_-dependent processes, while the lack of B_12_ production at human body temperatures creates avenues for therapeutics interfering with B_12_ supply. Altogether, this work highlights a gradual rather than stepwise reprogramming of the *P. aeruginosa* proteome in response to environmental cues, and highlights proteomics as a comprehensive tool to investigate system level mechanisms of challenging pathogens.

**Author Summary:** *Pseudomonas aeruginosa* is a bacterial pathogen that causes severe infections in human patients. This organism restructures its cellular machinery significantly between the low temperatures in the environment and the high body temperatures of a human host. Several factors influence its virulence and pathogenicity, such as the supply of essential metals (iron, zinc, cobalt) and vitamins as bacterial nutrients. However, how the availability of cobalt, together with varying temperature, regulates *Pseudomonas aeruginosa* remains poorly understood. Here, we applied measurements of proteomics and cobalamin, a cobalt-containing essential vitamin, to understand the changes of this pathogen in response to changing environmental conditions. Our study reveals that this bacterium reacts to varying temperatures by continuously modulating biofilm formation, bacterial movement, cellular metabolism, and virulence expression. We also found that *Pseudomonas aeruginosa*’s capacity to produce its own cobalamin declines at host-associated temperatures, while the vitamin’s concentration is kept constant within the cell at all temperatures. These outcomes are important because starving this pathogen of cobalamin could present a novel therapeutic strategy for combating *Pseudomonas aeruginosa* infections.

## Introduction

*Pseudomonas aeruginosa* is a ubiquitous high-priority pathogen that poses a major global health threat.[1] Its ability to persist in a wide range of settings, from medical facilities to urban soils and ocean waters, creates a pervasive exposure risk for humans and a widespread source of initial infections.[2] *P. aeruginosa* can cause acute or chronic sepsis, pneumonia, enterocolitis, and dermatitis, as well as opportunistic cross-infections of existing conditions.[3] Due to its increasing prevalence of resistance against multiple antibiotics and host defense strategies, it is especially dangerous in nosocomial infections of immunocompromised patients.[1, 4, 5]

The pathogenicity of *P. aeruginosa* is closely linked to its ability to form biofilms and to tolerate suboxia, as well as to external cues such as temperature and metal availability. *P. aeruginosa* biofilms – sessile multicellular communities that attach to surfaces through a cohesive matrix built from extracellular polymeric substances (EPS) – are responsible for the majority of human infections due to enhanced multidrug resistance and defense protection.[6] The transitions between planktonic twitching, swimming, or swarming motility and biofilms are governed by complex gene regulation mechanisms that dynamically respond to environmental stimuli, e.g. through two-component sensing systems,[7, 8] and induce quorum sensing circuits which further advance biofilm maturation.[9]

Its capabilities for anaerobic growth with various electron acceptors (i.e. nitrate) enable this pathogen to grow along steep oxygen gradients,[10] thus further enhancing its virulence and persistence. *P. aeruginosa* can colonize local hypoxic zones in chronic wounds and cystic fibrosis mucus of the host organism, and exacerbate them through oxygen consumption and anaerobiosis in the deeper layers of biofilms.[11]

External temperature is another important cue for bacterial host colonialization. Indeed, the switch from ambient (∼22 °C) to host-associated (∼37 °C) temperatures does not only shape the structural integrity of biofilm architecture,[6] but also dictates other adaptations of *P. aeruginosa* to the human body through, for instance, thermoregulated RNA folding.[12] Further temperature-induced changes include the upregulation of bacterial secretion systems, the expression of virulence factors and quorum sensing molecules, the acquisition of essential nutrients, and the speed of metabolism and multiplication.[12–16] While 37 °C is commonly regarded as host condition, the lower temperatures of wound beds (30-33 °C)[17] and the upper trachea (35.5 °C)[18] suggests that *P. aeruginosa* may adapt to a wider range of temperatures during infection.

Metal ion supply and homeostasis are other crucial determinants for pathogenicity. Iron (Fe), for example, is essential for biofilm formation, increases antibiotic resistance, and contributes to host tissue damage within the secreted compound pyocyanin.[19] In fact, nutritional immunity is a defense strategy of the human host to withhold Fe and other metals from infecting bacteria,[20] inducing *P. aeruginosa* competition for Fe through siderophore expression and heme degradation.[21–23] In addition, Zinc (Zn) is necessary in metalloenzymes that function as virulence agents, such as tissue degrading proteases,[24, 25] and new insights about Zn and manganese (Mn) in host nutritional immunity against *P. aeruginosa* are starting to emerge.[26, 27]

In contrast, the roles of cobalt (Co) and the Co-containing vitamin B_12_ at the host-microbe interface during *P. aeruginosa* infections remain poorly understood. Co is an essential cofactor in isoforms of methionine synthetase, ribonucleotide reductase (RNR), and other metabolic enzymes.[28] Although *P. aeruginosa* only encodes genes for aerobic B_12_ biosynthesis, the pathogen requires B_12_ for both anaerobic biofilm formation[29] and host infection,[30] and the mechanisms of oxygen-devoid B_12_ homeostasis are still unclear.[10, 31] Interestingly, Co can partially compensate for Zn in starved *P. aeruginosa* and restore some virulence traits,[32] further demonstrating the need to understand the function of Co in human infections.

Global high-resolution proteomics is a powerful tool to investigate the functional status of pathogens under physiologically relevant conditions. Understanding *P. aeruginosa* gene regulation under varying temperatures has been identified as knowledge gap relevant to infection control.[33] To date, most system analyses of *P. aeruginosa* have relied on RNA-based approaches, often comparing two discrete growth temperatures (low vs. high),[12–15, 34] or focused on metal biology without accounting for temperature.[25, 35–37] Furthermore, several of these reports rely on older microarray techniques with reduced quantification accuracy compared to modern technologies.[13] Here, we apply global proteomics across a high-resolution temperature gradient to illuminate how metal metabolism and pathogenicity are reshaped across environment-to-host transitions in *P. aeruginosa*, and to investigate various metal and vitamin homeostasis mechanisms across a high-resolution temperature gradient.

## Methods

### Strain Cultivation

*Pseudomonas aeruginosa* 2-54 (PA254) was isolated in the Central Pacific Ocean on the research expedition METZYME (KM1128) aboard the R/V Kilo Moana, as described previously.[10] It is unclear if this new strain is of environmental origin or a human-derived contaminant. Due to its isolation from a marine setting, however, PA254 was cultured in autoclaved marine broth containing 1 L 0.2 µm-filtered coastal seawater (Vineyard Sound), 5 g peptone (Fisher Scientific), and 1 g yeast extract (BD Difco). Solid media were prepared with addition of 1.5% agar (Fisher Scientific).

### Growth Curves

To record growth curves at different temperatures (22, 25, 30, 35, 37, 40, 42 °C), PA254 was grown in 12-well plates in a SpectraMax M3 plate reader (Molecular Devices). First, an acclimated pre-culture of PA254 in 5 mL medium was prepared from a single colony or glycerol stock, and shaken at 600 rpm at the desired temperature overnight. Before inoculation of the 12-well plate, 20 mL medium, supplemented with 1 µM CoCl_2_, as well as the plate reader chamber were pre-adjusted to the desired temperature for at least 15 min. After that, 1.5 mL medium and 15 µL PA254 pre-culture were dispensed into each well. Optical density was then recorded at 600 nm every 10 min for 16-42 hours until reaching stationary phase, with orbital shaking for 5 s before every measurement. To prevent condensation, plate lids were pre-treated with 5% Triton-X in ethanol, and dried in a laminar flow hood overnight.

### Genome Annotation

Genomes of bacterial isolates from the METZYME expedition were sequenced at the Johns Hopkins Deep Sequencing and Microarray Core Facility, as reported previously.[10] Isolates 1-54 and 2-54 were identified to be the same strain of *P. aeruginosa*, and PA154 yielded a circular chromosome (6,455,702 base pairs). JSpeciesWS tetra correlation search (TCS)[38] showed P154 closely related to *P. aeruginosa* P-14 and BWH047 with a tetranucleotide-signature correlation score of Z = 0.99984; however, whole-genome alignments with these hits suggested a distinct and novel strain. Translation initiation sites were predicted with the gene finding software Prodigal (PROkaryotic DynamIc programming Genefinding Algorithm).[39] Open reading frames (6,523) were then translated into amino-acid code, and a Diamond-search was performed against the NCBI nr (non-redundant) protein database, resulting in 5,752 functionally annotated and 793 hypothetical proteins. Of the 793 hypothetical proteins, 269 could be annotated using eggNOG (evolutionary genealogy of genes: Non-supervised Orthologous Groups) mapper[40] and were integrated into the final annotated proteome.

### Proteomics

All solvents utilized here were LC-MS grade (Fisher Optima), unless otherwise noted.

#### Extraction

Protein extraction and digestion were performed using a modified version of the Protifi S-trap mini spin column protocol recommended by the manufacturer (https://protifi.com/protocols). Briefly, 9 mL of a PA254 culture were harvested in a 2 mL EtOH-washed microtube through repeated centrifugation at 9,600 × g for 5 min, and frozen at −80 °C prior to extraction. Upon thawing, the pellet was resuspended in 300 µL lysis buffer (2% SDS, 50 mM tetraethylammonium bromide, pH 8.5) and incubated at 95 °C for 10 min. The samples were cooled on ice, treated with 2 µL benzonase nuclease (25.5 u/µL, Novagen), shaken at 350 × g for 30 min at 37 °C, and centrifuged at 9,600 × g for 10 min. The supernatant was transferred to a fresh 2 mL EtOH-washed microtube and total protein quantified via micro-BCA protein assay (Thermo Scientific).

#### Reduction and Alkylation

An aliquot of 100 µg extracted proteins in 400 µL lysis buffer (2% SDS, 50 mM tetraethylammonium bromide, pH 8.5) was prepared for reduction and alkylation. First, 4 µL 500 mM dithiothreitol (in 50 mM ammonium bicarbonate) were added, and samples incubated at 45 °C for 30 min. Then, 12 µL 500 mM iodoacetamide (in 50 mM ammonium bicarbonate) were added, and samples incubated at room temperature for 30 min in the dark. Excess iodoacetamide was quenched by addition of 4 µL 500 mM dithiothreitol (in 50 mM ammonium bicarbonate). Afterwards, samples were treated with 23 µL 12% phosphoric acid, incubated at room temperature for 5 min, and diluted with 1.7 mL binding buffer (100 mM tetraethylammonium bromide, pH 7.1, 90% MeOH). Proteins were loaded onto pre-rinsed S-trap mini-spin columns in increments of 600 µL at 1,100 × g for 30 s, with each flow-through being loaded a second time. The samples were washed with 8-10 times with 600 µL binding buffer (100 mM tetraethylammonium bromide, pH 7.1, 90% MeOH) at 1,100 × g for 30 s, and one time with 600 µL 90% MeOH at 9,600 × g for 2 min.

#### Digestion

Trypsin digestion was performed on-column with a protein:trypsin ratio of 25:1. A solution of 4 µL trypsin (Trypsin Gold, Mass Spectrometry Grade, Promega) in 125 µL 50 mM ammonium bicarbonate was allowed to permeate the S-trap mini spin columns by centrifuging at 70 × g for 30 seconds, and reloading any flow-through. The digestion was carried out at 37 °C for 14 hours.

#### Peptide preparation

Digested proteins were eluted into a fresh 2 mL EtOH-washed microtube stepwise with 80 µL 50 mM Ammonium bicarbonate, 80 µL 0.2% formic acid, and 80 µL 50% acetonitrile and 0.2% formic acid at 9,600 × g for 1-5 min each. Combined eluents were then quantified via micro-BCA protein assay (Thermo Scientific) before drying in a SpeedVac (ThermoSavant) at room temperature. The residue was redissolved in LC-MS buffer (2% acetonitrile, 0.1% formic acid) to a final concentration of 1 µg/µL peptides. To avoid particulates, peptides were centrifuged at 9,600 × g for 20 min, and the supernatant diluted to 0.1 µg/µL for analysis.

#### Peptide analysis

Tryptic peptides were analyzed using liquid chromatography coupled with tandem mass spectrometry (LC-MS/MS) by injecting 1 µg on a Thermo Vanquish Neo HPLC reverse phase chromatography system paired with a Thermo Scientific Orbitrap-Astral mass spectrometer using a Thermo Flex nanospray source. Each sample was concentrated onto a trap column (0.3 × 10 mm ID, 5 µm particle size, 100 Å pore size, C18 Thermo PEPMAP100) and rinsed with 100 µL 0.1% formic acid in water before gradient elution through phase C18 column (0.1 x 500 mm ID, 2 µm particle size, 120 Å pore size, C18 Reprosil-Gold, Dr. Maisch GmbH) using a 500 nL/min flow rate. Liquid chromatography was performed using a nonlinear 60-minute gradient from 1% to 95% buffer B (0.1% formic acid in 80% acetonitrile) in buffer A (0.1% formic acid in water). Data dependent acquisition (DDA) was used with a precursor range of 380 m/z to 1280 m/z on the Orbitrap at 240 K resolution. MS2 scans were performed with 1.6 m/z MS2 windows on the Astral analyzer with a range of 150 2000 m/z, 2-6 m/z charge states, a dynamic exclusion of 7 sec, and a 2 sec cycle time between master scans.

### Proteomics Informatics

Raw mass spectra were searched against the PA154 annotated proteome fasta, generated as described under ‘genome annotation’, using the program FragPipe (Version 23). The parameters used were a precursor mass tolerance of 5 ppm, a fragment mass tolerance of 10 ppm, and up to 2 missed cleavages. Differential expression analysis was conducted using FragPipe-Analyst[41] based on limma (Linear Models for Microarray and Omics Data). MaxLFQ intensity values were processed using a Benjamini Hochberg adjusted *p*-value < 0.05, a log2 fold-change cutoff of 1, and no imputation, to yield overexpressed (positive), underexpressed (negative), and differentially expressed (total) proteins (**Dataset S1**). Volcano plots of differently expressed proteins were prepared by plotting the -log10 unadjusted *p*-value against log2 fold-change using Python. For visualization purposes, a reduced log2 fold-change cutoff of 0.5 was used. Heatmap plots of proteins versus temperature were prepared in the following manner: ANOVA was applied to test for the dependence of protein abundance on temperature using three biological replicates at each temperature point. Proteins where ANOVA *p* < 0.05 were visualized as heatmaps using R studio using unique spectral counts max-normalized to a scale of 0-1.

### Pathway Analysis

For metabolic pathway analysis, overrepresentation analysis (ORA), and gene set enrichment analysis (GSEA), the predicted proteome of PA254 was first mapped onto Kyoto Encyclopedia of Genes and Genomes (KEGG) pathway maps. For this purpose, the KEGG pathway maps of the *P. aeruginosa* PAO1 proteome were used, which are publicly available (https://www.kegg.jp/kegg-bin/get_htext?pae00001). A Diamond search was applied to assign known PAO1 pathways to the corresponding PA254 proteins, leading to 6,036 proteins with pathways. For the remaining 487 proteins without pathways, additional pathway information for 197 proteins was obtained using eggNOG mapper[40] and integrated into the final pathway database (*Supplemental Dataset S2*). Differentially expressed proteins per pathway per temperature were then plotted as percent of total detected proteins per pathway per temperature.

Overrepresentation analysis (ORA) was performed on the over- and underexpressed proteins of different temperatures vs. 22 °C (foreground), obtained as described under ‘proteomics informatics’, against total detected proteins per pathway per temperature comparison (background). Using Python, the statistical significance of pathway overrepresentation was assessed using Fisher’s exact test with a one-sided alternative hypothesis, and *p*-values were corrected for multiple testing using the Benjamini–Hochberg false discovery rate (FDR) method. Pathways with an FDR < 0.05 were considered significantly overrepresented, and plotted against -log10 FDR.

Gene set enrichment analysis (GSEA) was performed on the total expressed proteins of different temperatures, obtained as described under ‘proteomics informatics’, by ranking all log2 fold-changes for each temperature comparison. Using Python, pathways for PA254 proteins were then converted into a Gene Matrix Transposed (GMT) format. GSEA was conducted using the package *gseapy* with 1,000 permutations per analysis. Pathways containing < 3 or > 500 proteins were excluded. Pathways with an FDR < 0.05 were considered significantly enriched, and plotted against the normalized enrichment score (NES).

### Cellular Cobalamin

All solvents were LC-MS grade (Fisher Optima), unless otherwise noted. To quantify the cellular cobalamin content, 9 mL of a PA254 culture were harvested in a 2 mL microtube through repeated centrifugation at 9,600 × g for 5 min. Cell pellets were washed twice with 1 mL 0.2 µm-filtered and autoclaved coastal seawater (Vineyard Sound) to remove dissolved extracellular cobalamin, and frozen at −80 °C prior to extraction. Upon thawing, samples were treated with 1.5 mL MeOH and 10 µL 10% HCl (for roughly 10 mM or pH 2 final concentration), and shaken at 800 × g for 30 min at room temperature in the dark. The supernatant was collected through centrifuging at 9,600 × g for 2 min, and the pellet extracted a second time as above. The combined extraction fractions were then dried in a SpeedVac (ThermoSavant) at room temperature in the dark before resuspension in 50 µL LC-MS buffer (2% acetonitrile, 0.1% formic acid). To avoid particulates, samples were centrifuged at 13,100 × g for 20 min in the dark, transferred to a 0.2 mL microtube, and centrifuged again. The supernatant was transferred to an amber microvial and measured via LC-MS/MS on a Thermo Dionex NC-3500RS HPLC coupled to a Thermo Scientific Fusion Orbitrap mass spectrometer with a Thermo Flex source. Each sample was concentrated onto a trap column (0.2 × 10 mm ID, 5 μm particle size, 120 Å pore size, C18 Reprosil-Gold, Dr. Maisch GmbH) and rinsed with 100 µL LC-MS buffer (2% acetonitrile, 0.1% formic acid) before elution through a reverse phase C18 column (0.1 x 150 mm ID, 3 μm particle size, 120 Å pore size, C18 Reprosil-Gold, Dr. Maisch GmbH). Chromatography was performed at 0.5 mL/min flow rate over a 40 minute nonlinear gradient from 5% to 95% acetonitrile and 0.1% formic acid in water. Mass spectrometry targeted the observable masses of cyanocobalamin (CN-B_12_; m/z = 1354.57, 678.29), methylcobalamin (Me-B_12_; m/z = 1343.59, 672.80), and hydroxocobalamin (OH-B_12_; m/z = 1345.57, 673.79). Adenosylcobalamin was not included in this study, as it was expected to be a minor contributor. Total MS2 fragment areas for each form of cobalamin (CN, Me, OH) were obtained using Skyline (Version 23.1) and used to calculate absolute concentrations. A 6-point calibration curve for each cobalamin (CN, Me, OH) was prepared in the range of 1-100 µg/L in LC-MS buffer (2% acetonitrile, 0.1% formic acid). External standards containing 5 µg/L of each cobalamin (CN, Me, OH) were analyzed with each set of unknown samples. A series of blanks was run after each standard or sample, and cobalamin peak areas eluting in blanks (if any) were added to the standard or sample peak areas until blanks washed out clean.

### Dissolved Cobalamin

Extracellular dissolved cobalamin in the spent media of PA254 cultures was measured according to a previously published method.[42] Due to representing only a small fraction (< 10 %) of total cobalamin, dissolved cobalamin was omitted from the discussion.

### Data Availability

The sequenced *P. aeruginosa* genome for the isolate 1-54 is available at Zenodo (doi: 10.5281/zenodo.15336754) and NCBI (BioProject PRJNA1541303). The annotated proteome has been submitted to PRIDE and NCBI. The raw global proteomic spectra have been submitted to ProteomeXchange and PRIDE (review access available upon request). The processed global proteomic data are available as **Dataset S1**.

## Results

### Growth at temperatures

In order to investigate the dependence of metal homeostasis and pathogenicity in *P. aeruginosa* PA254 on external temperature, the organism was cultivated across a range from 22-42 °C, in increments of 2-5 °C. (**Fig 1**). Pre-cultures acclimated to each desired temperature overnight were used for inoculation. PA254 was capable of growing to a dense culture (OD_600_ > 0.35) in all conditions tested, albeit with a longer lag phase at lower temperatures and a lower maximum OD_600_ at higher temperatures (**Fig 1a**). The growth rates continuously increased from 0.10 hr^−1^ at 22 °C to 0.45 hr^−1^ at 42 °C (**Fig 1b**).

**Fig 1.**
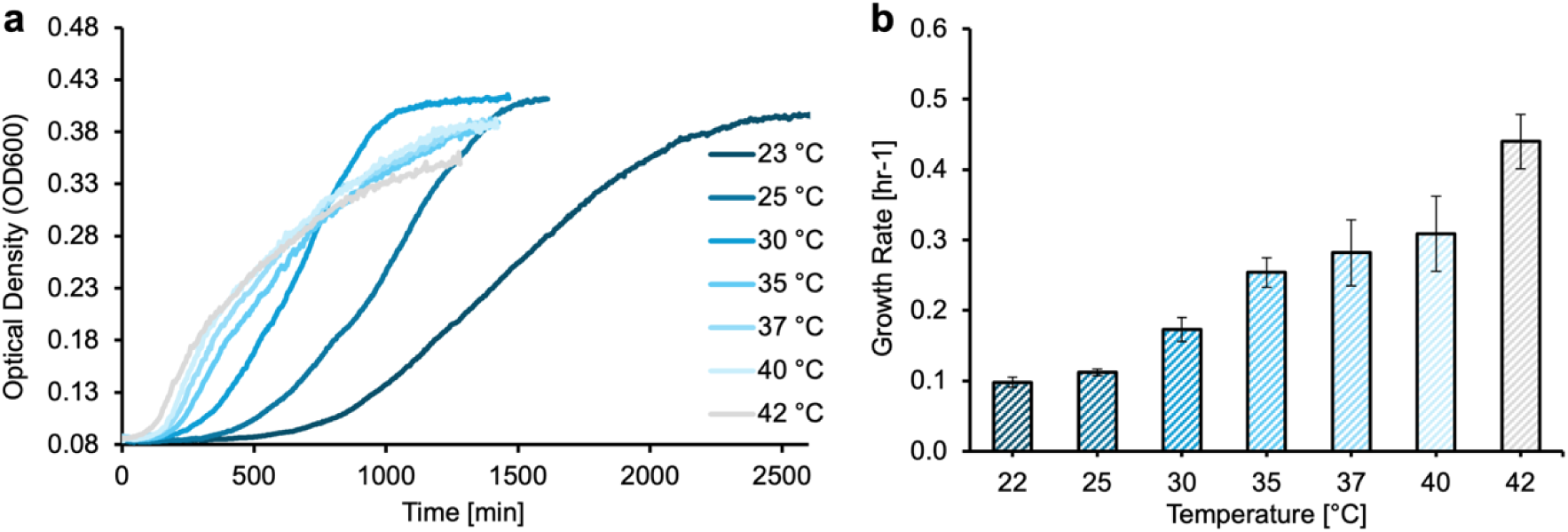
Temperature-dependent average growth curves (a) and growth rates (b) of *P. aeruginosa* PA254 cultivated at environmental (22-30 °C), host-associated (35-37 °C), and heat stress conditions (40-42 °C). Error bars represent one standard error of the mean, *n* = 2 or 3. For clarity, standard deviations for (a) are displayed in **Fig S1.**

### Global proteomics metrics

Cultures in stationary phase were evaluated through global proteomics analyses. Using our sequenced genome for peptide mapping, we identified 4,730 unique proteins in this study, representing a proteome coverage of 72.5 % across all temperatures (**Dataset S1**). Individual samples were yielded 3,400-4,700 unique proteins (**Fig S2**). The coefficient of variation, pairwise Pearson correlation, and log2 centered intensity showed strong reproducibility within replicates (**Figs S3-S5**). Principal component analysis revealed two components with 34.8% (PC1) and 21.1% (PC2), indicating that experimental treatments accounted for the majority of observed variance (**Fig S6**). Up to 13.5 % of all detected proteins were differentially expressed (**Fig S7**), with the largest amount between the largest temperature differential of 22 °C vs. 42 °C (638 proteins; **Fig S8**), and the smallest amounts between 25 °C vs. 30 °C, 30 °C vs. 35 °C, and 37 °C vs. 40 °C (< 3 proteins).

### Differential protein expression

Major metabolic and physiological traits known to be temperature-responsive in *P. aeruginosa* were reflected by significant changes in the proteome (ANOVA *p* < 0.05; **Fig 2**). For instance, classic heat stress proteins increased with temperature and reached their maxima at 42 °C, including the molecular chaperones ClpB, GroEL, DnaJ, and DnaK (**Fig 2a**), similar to the heat shock response of other Pseudomonads.[15, 43]

**Fig 2.**
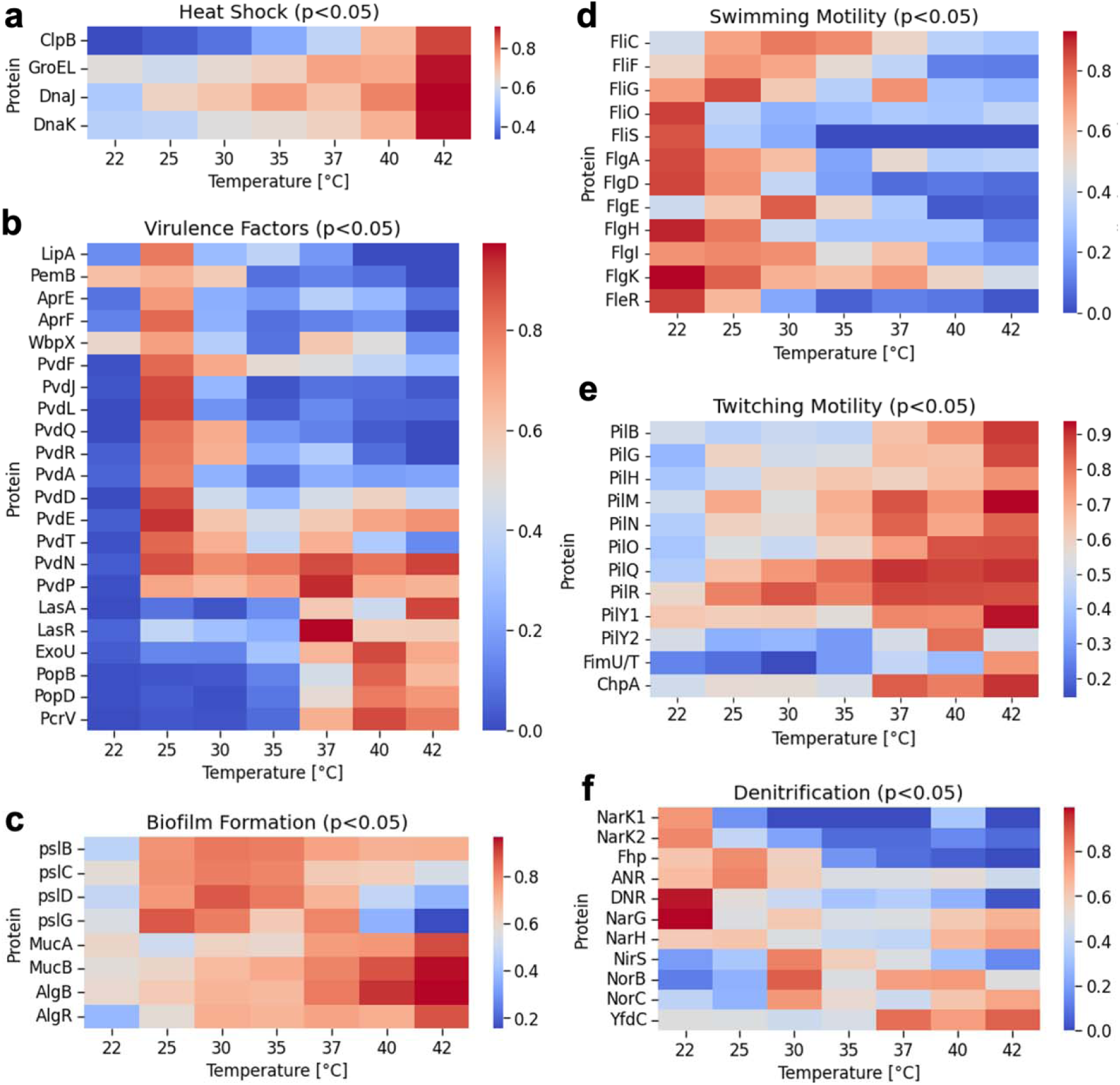
Temperature-dependent proteome shifts related to virulence (a), heat shock (b), biofilm formation (c), twitching and swarming motility (d), swimming motility (e), and denitrification (f) in *P. aeruginosa* PA254. Displayed are only entries that showed a significant correlation of protein abundance to temperature gradient (ANOVA *p* < 0.05), and that were differentially abundant between at least two temperature points (Benjamini-Hochberg adjusted *p* < 0.05). For abbreviations, see *Table S1*.

PA254 expression of several established virulence factors was highly temperature dependent and partitioned into two groups (**Fig 2b**). One set of proteins showed maximum protein abundances at 25 °C, and contained the sugar antigen producing rhamnosyltransferase WbpX,[44] several members of the pyoverdine gene cluster Pvd,[45] as well as effectors released by type 1-3 bacterial secretion systems (T1SS-T3SS),[46] namely the surface structures AprE and AprF (T1SS), the exotoxin LipA (T2SS), and the secreted product PemB (T3SS). The second set of proteins was more abundant at 37-42 °C and included the secreted tissue-degrading protease LasA and corresponding regulator LasR, and select effectors of T3SS, such as exotoxin ExoU, the pore-forming proteins PopB/D, and the needle tip protein PcrV.[46]

Proteins implicated in biofilm formation also formed two temperature response groups according to initial and mature biofilm processes. The EPS biofilm matrix of *P. aeruginosa* largely consists of exopolysaccharides synthesized via the Psl operon, which was most abundant between 25-35 °C (**Fig 2c**), and is associated with early cell aggregation and biofilm initiation.[47] Alginate, on the other hand, is a major constituent of chronic mucoid biofilms; AlgB/R upregulate the AlgACD gene locus which directly controls the biosynthesis of alginate precursors.[47] Both AlgB/R, and the negative regulators MucA/B protecting *P. aeruginosa* from alginate overproduction,[47] were most abundant at 37-42 °C.

Bacterial swimming motility operates via rotating flagella, where the individual components are constructed using the large gene clusters Fli (rotor, filament and cap) and Flg (hooks, rings and rod). Many of these proteins were highly abundant below 30 °C, including the flagellin building block FliC (**Fig 2d**). The response regulator FleR is additionally implicated in swarming, a behavior connected to initial biofilm development and antibiotic resistance,[48] and had a significantly higher abundance at 22-25 °C (**Fig 2d**).[49] Bacterial twitching motility, on the other hand, is a flagella-independent movement along moist, solid surfaces with the help of hair-like pili. More than ten proteins of the associated Pil operon were of higher abundance above 37 °C, such as the minor pili assembly subunit protein FimUT, and the twitching sensor kinase ChpA (**Fig 2f**).[50]

Under anaerobic conditions, *P. aeruginosa* operates via denitrification using alternative electron acceptor for energy production. Despite aerobic growth here, denitrification proteins occurred across the temperature gradient, albeit at relatively low spectral counts (**Dataset S1**), and exhibited diverse trends (**Fig 2g**). Several nitrate/nitrite transporters (NarK), nitrate respiration regulators (ANR, DNR), and nitrate reductases (NarGH) were most abundant in the 22-25 °C range. A nitrite reductase (NirS) and nitric oxide reductases (NorBC) peaked at 30 °C, and the nitrite/formate transporter YfdC at 37-42 °C. However, select proteins (NarG/H, NorB/C) were expressed at two or more non-contiguous temperature regimes.

### Pathway analyses

To identify functional programs of *P. aeruginosa* responding to temperature stimuli, proteins were mapped to the corresponding cellular pathways of the Kyoto Encyclopedia for Genes and Genomes (KEGG) Pathway Database (*Supplemental Dataset S2*). A comparison between the differentially expressed proteins at temperature minimum vs. maximum (22 °C vs. 42 °C) showed that changes concentrated mostly within the pathways of xenobiotics degradation (16.5 %), cell motility (15.7 %), lipid metabolism (14.8 %), and bacterial infectivity (14.3 %; **Fig S9**).

Both overrepresentation analysis (ORA) and gene set enrichment analysis (GSEA) were applied to identify statistically significant pathway trends (**Fig 3**). ORA detects pathways based on differentially expressed proteins, while GSEA captures broad trends across the entire ranked proteome. ORA showed an enrichment in a subset of 23 and GSEA of 24 metabolic pathways, with partial overlap.

**Fig 3.**
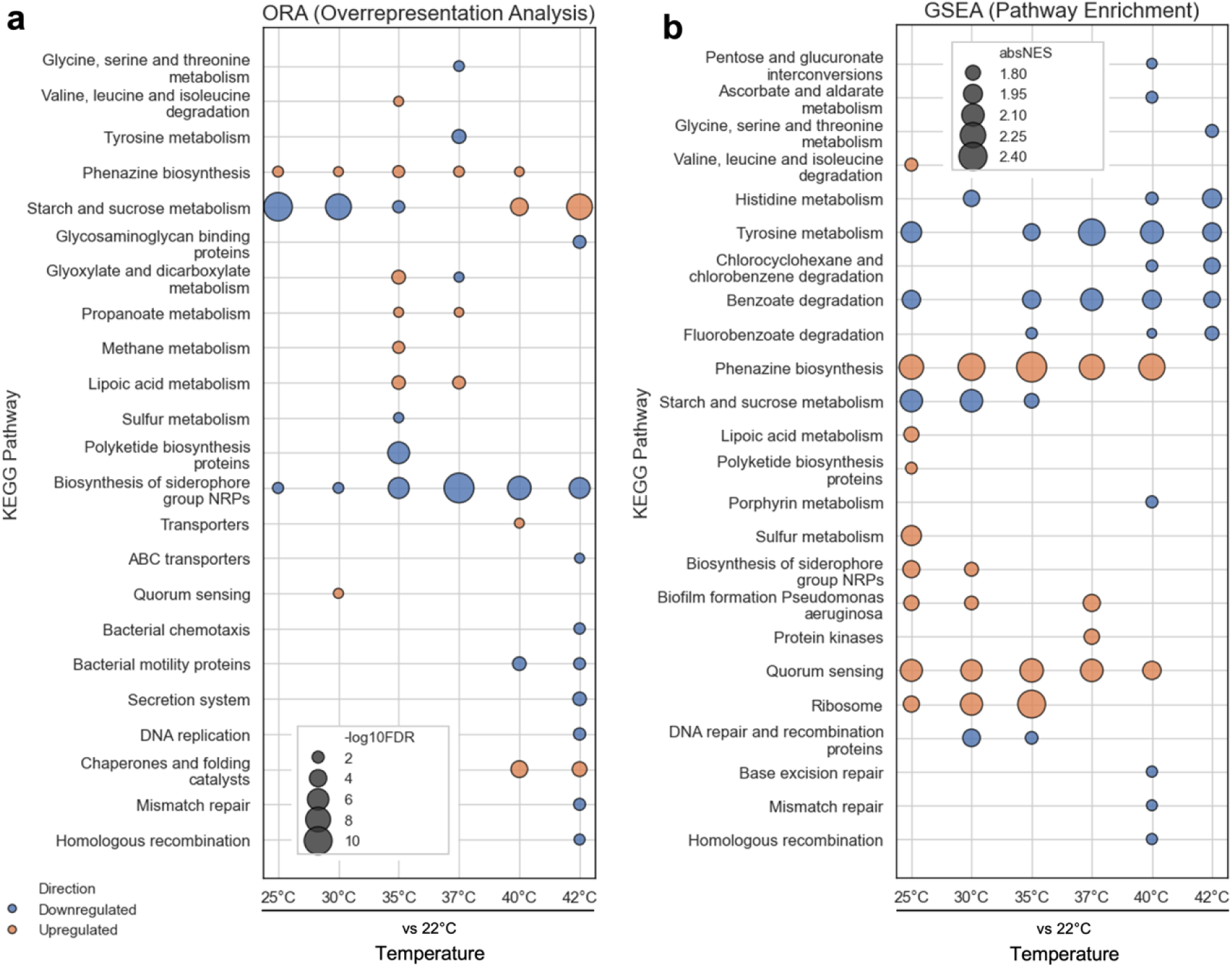
Temperature-dependent pathway analysis of *P. aeruginosa* PA254 via a) ORA (overrepresentation analysis) and b) GSEA (gene set enrichment analysis). Less abundant proteins at temperatures relative to 22 °C displayed in *blue*, and more abundant proteins in *orange*. For ORA, proteins with a Benjamini-Hochberg adjusted *p* < 0.05 are displayed as -log10 FDR (false discovery rate). For GSEA, proteins with an FDR *q*-value < 0.05 are displayed as absolute NES (normalized enrichment score).

Several coordinated metabolic shifts were captured with both ORA and GSEA approaches. The metabolism of certain amino acids was negatively enriched at temperatures above 37 °C, e.g. glycine, serine, threonine, histidine, and tyrosine (**Fig 3 a vs. b**). Biosynthesis of the virulent pigment phenazine showed strong, coherent enrichment between 25-40 °C compared to both temperature extremes (22 or 42 °C). Quorum sensing proteins were similarly enriched throughout intermediate temperatures but not at extremes. Starch and sucrose metabolism were negatively enriched below 37 °C and positively enriched above 37 °C, mirroring the trend in growth rate (**Fig 1b**). Under prolonged heat stress conditions (40–42 °C), genetic information processing pathways started to decline, including DNA replication, base excision repair, mismatch repair, and homologous recombination.

ORA further highlighted an overrepresentation of carbon assimilation and energy production pathways at host-associated temperatures (35–37 °C), namely glyoxylate, propanoate, methane, and lipoic acid metabolism. Heat stress (40–42 °C) resulted in a decline in bacterial chemotaxis and motility pathways, while chaperone and folding catalysts were overrepresented (**Fig 3a**).

GSEA revealed a negative enrichment in the catabolism of complex carbohydrates (pentose and glucuronate interconversion, ascorbate and aldarate metabolism) at high temperatures (40–42 °C). In addition, catabolic pathways for xenobiotics (chlorocyclohexane, chlorobenzene, benzoate, and fluorobenzoate degradation) also showed a concerted negative enrichment and temperatures above 35 °C. Ribosomal proteins were continuously enriched between 25–35 °C (**Fig 3b**).

A small subset of pathways exhibited contrasting trends between ORA and GSEA. The biosynthesis of polyketides and non-ribosomal peptides (NRPs) was negatively enriched in ORA, but positively enriched at lower temperatures using GSEA (25–30 °C); these secondary metabolites include antibiotics and siderophores.[45] This usually happens when distinct or unequally sized subsets of one pathway are captured by each different method.[51]

### Metal homeostasis

Iron uptake systems in *P. aeruginosa* are directly correlated with host infection,[19–23] and 13 proteins associated with these mechanisms changed significantly with temperature in this study (ANOVA *p* < 0.05, **Fig 4a**). Among these, alternative iron uptake systems[52] were significantly more abundant above 37 °C when compared to 22 °C: the outer membrane heme receptor PhuR, the hemopexin-binding importer HxuA, and the ferri-enterobactin transporter PirA. FemA, responsible for ferri-mycobactin uptake, and FoxA and FiuA, both involved in ferrichrome/ferrioxamine uptake, were significantly less abundant at 22-30 °C when compared to 42 °C. In addition, the abundance of FoxA was also significantly increased at 37-42 °C when compared to 25 °C. The FpvAB system transports Fe-bound pyoverdine and was significantly elevated at all temperatures above 22 °C.

**Fig 4.**
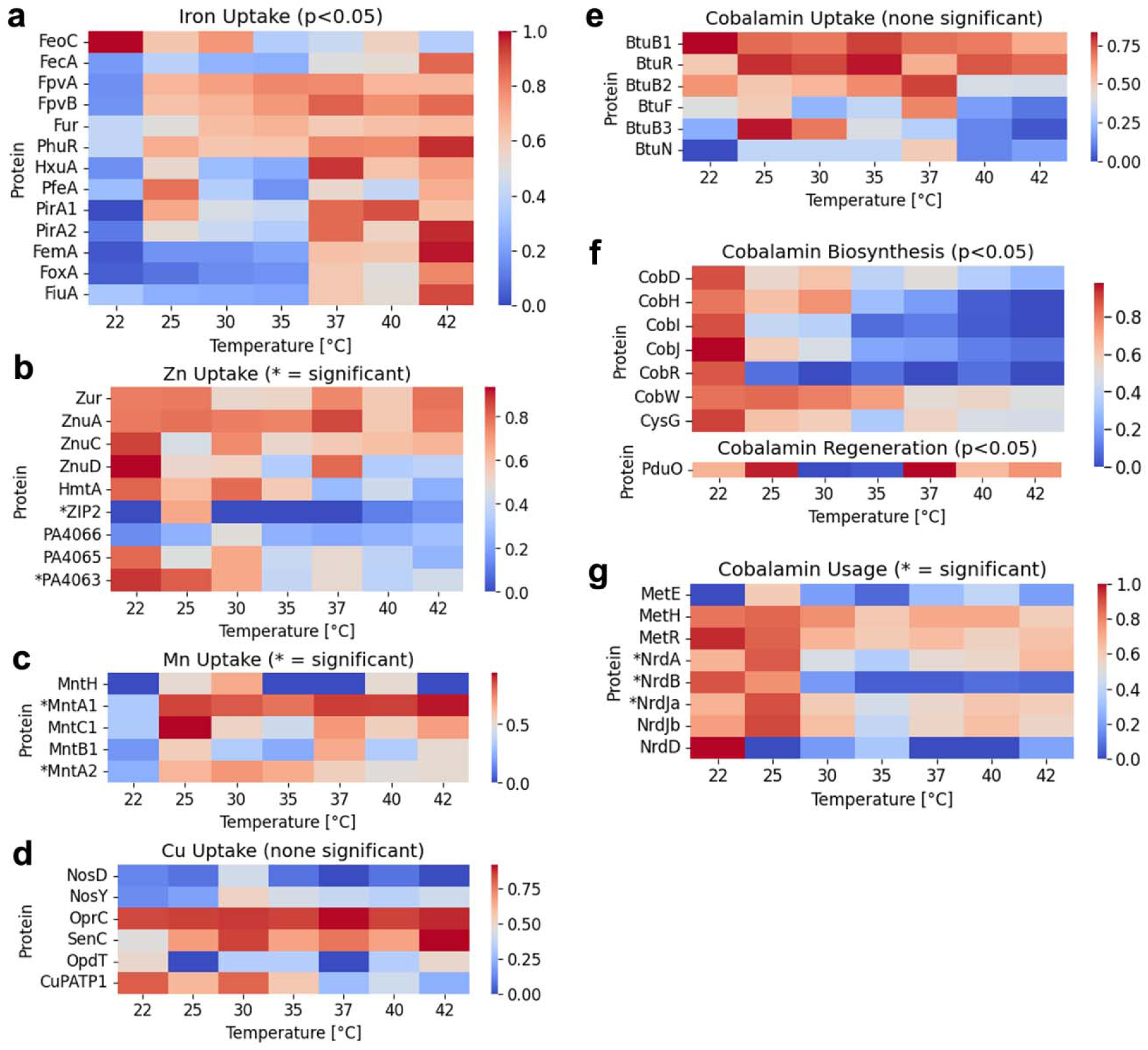
Temperature-dependent metal homeostasis in *P. aeruginosa* PA254. Changes in protein abundance related to Fe uptake (**a**), Zn uptake (**b**), Mn uptake (**c**), Cu uptake (**d**), cobalamin uptake (**e**), cobalamin biosynthesis and regeneration (**f**), and cobalamin usage (**g**). Only entries that showed a significant correlation of protein abundance to temperature gradient (ANOVA *p* < 0.05), and that were differentially expressed between at least two temperature points (Benjamini-Hochberg adjusted *p* < 0.05), are displayed. For gene abbreviations, see *Supplemental Material Table S1*.

Pseudomonads possess multiple redundant uptake strategies for other biologically relevant metal ions and their complexes (Co, Zn, Mn, copper (Cu)).[25, 52] Only two Zn transporters[25] and one Mn uptake protein[53] varied significantly with temperature, but were not differentially abundant at host-adapted conditions (**Fig 4b-c**). No significant responses to host temperatures were found for Cu uptake systems (**Fig 4d**). While no Co metal ion-specific transporter in *P. aeruginosa* is known, Co is often co-imported through other channels (e.g., with Zn), and cobalamin imported through TonB-dependent BtuB receptors.[54] Three putative B_12_ transporters (BtuB) were identified in the PA254 proteome; however, no Co or B_12_ import system was temperature-sensitive (**Fig 4e**). Proteins related to the Zn/Co siderophore pseudopaline[55] were not detected, consistent with its expression under Zn-limited conditions not included in this study.[25]

Besides extracellular cobalamin uptake, *P. aeruginosa* is capable of aerobic B_12_ biosynthesis through the Cob cluster. Several members of the biosynthetic Cob pathway as well as CysG varied strongly with temperature, and the abundance these proteins diminished significantly above 22 °C (**Fig 4f**). These responsive Cob genes stem from all major Cob gene neighborhoods.[54] CobA/F/L/ST/V were not observed in this dataset; CobC/G/K/M-O/Q/T/U did not change significantly. The corrinoid adenosyltransferase PduO catalyzing the regeneration of active cobalamin cofactors was highest at 37 °C, but not differentially expressed at that temperature (unadjusted *p* = 0.018).

Cobalamin is employed as cofactor for the methionine synthase MetH and the ribonucleotide reductase NrdJ. The cobalamin-free counterparts of these enzymes are MetE, NrdA/B under oxic conditions, and NrdD under anoxic conditions, respectively.[31] B_12_-dependent MetH was detected with no significant changes throughout the temperature gradient, along with the methionine synthesis regulator MetR (**Fig 4g**). Both B_12_-dependent and independent NrdA/D/J changed significantly with temperature with higher abundances at low temperatures versus 35 °C. NrdJa was also more abundant at 25 versus 40-42 °C. Consistent with the aerobic culturing conditions, the anoxic protein NrdD had very low spectral counts (**Fig S10**), and was not detected at 25 and 37-40 °C.

### Cobalamin concentrations

The absolute concentrations of cobalamin within *P. aeruginosa* PA254 cells were measured using LC-MS/MS. For that purpose, the amounts of the analogs cyanocobalamin (CN-B_12_), methylcobalamin (Me-B_12_), and hydroxocobalamin (OH-B_12_; **Fig S11**) were quantified individually and summed for total B_12_. To ensure detectable B_12_ concentrations, the growth medium was supplemented with 1 μM CoCl_2_ (**Fig 5a**). This medium-level addition also prevents cells from experiencing B_12_ excess or starvation.[56] Intracellular cobalamin through the temperature curve averaged at 0.384 ± 0.026 pM OD^− 1^ when normalized to optical density of the PA254 culture, and the values at individual temperatures were not statistically different (**Fig 5b**). The majority of observed cobalamin was OH-B_12_ at all temperatures (**Fig S12**). Extracellular cobalamin in the growth medium represented only a small fraction (< 10%) of the total B_12_ and was thus omitted from this study. It was found that PA254 produced B_12_ until late stationary phase, and that intracellular B_12_ did not decline within aged cultures (**Fig S13**).

**Fig 5.**
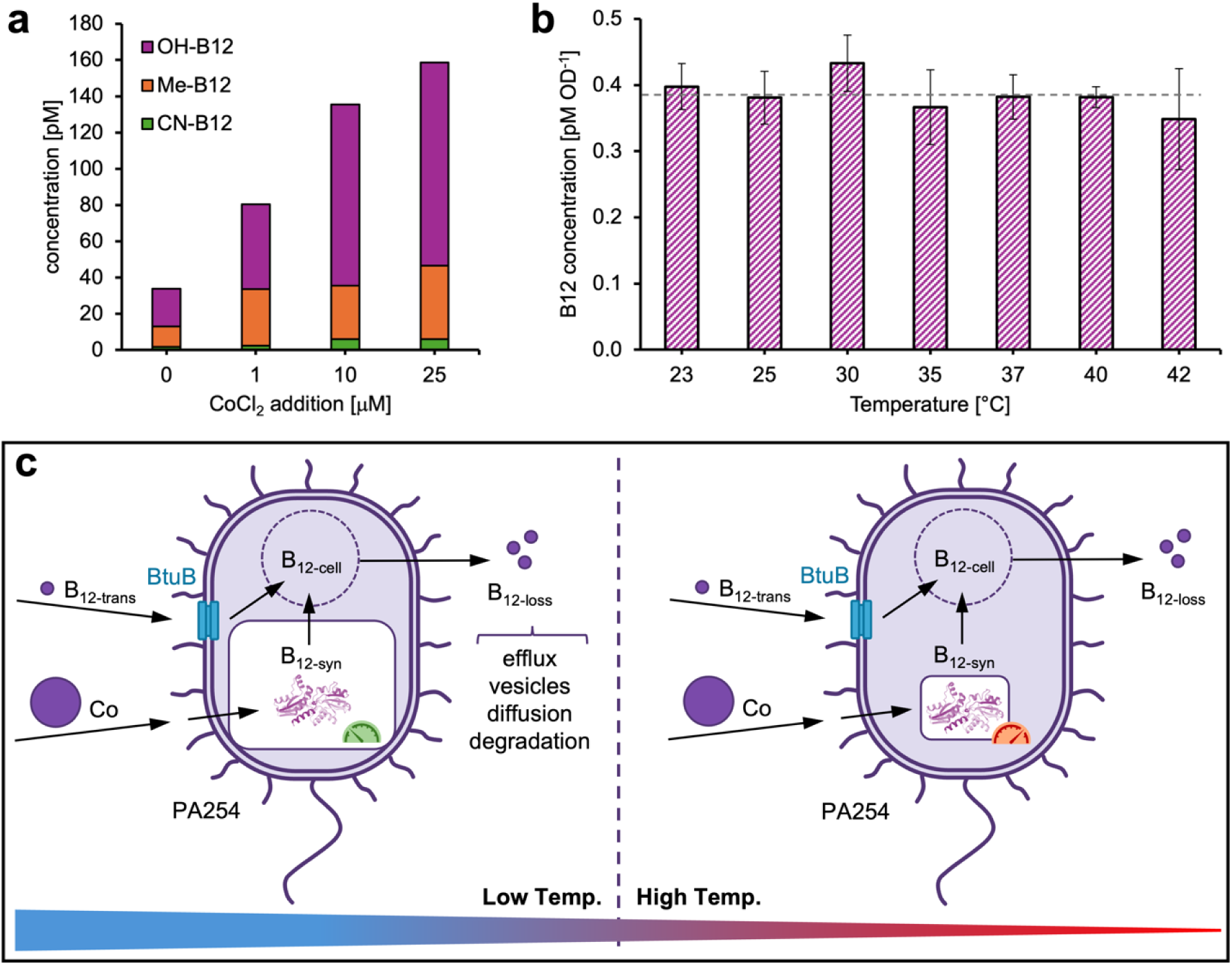
Metrics of cobalamin metabolism in *P. aeruginosa* PA254. **a)** Intracellular cobalamin concentrations in PA254 cultures supplemented with cobalt chloride. **b)** Intracellular cobalamin concentrations in PA254 across the temperature gradient, with dashed line denoting the average. **c)** Schematic representation of cellular B_12_ processes including uptake (B_12-trans_), biosynthesis (B_12-syn_), reservoir (B_12-cell_), loss (B_12-loss_), and enzyme efficiency (speedometer).

## Discussion

Infectious strains of *P. aeruginosa* have been reported to operate under two distinct temperature regimes: an environmental lifestyle (∼20-25 °C) favoring biofilm growth plus the degradation of complex natural sugars and xenobiotics, and at human body temperatures (37 °C) favoring rapid proliferation plus adaptation to nutrients scavenged during host infection.[6, 12, 14–16, 57, 58] Here, this study shows a continuous transition of these two modes over a temperature gradient, and associated adaptations in pathogenic traits, biofilm formation, metal homeostasis, and metabolism throughout.

### Continuous modes of temperature-dependent pathogenic physiology

Prior experiments with the virulent reference strain PAO1 established that many quorum sensing products, secretion systems, and virulence factors are strongly induced at 37 °C,[14] especially factors reliant on the transcriptional regulator *rhlR*[12] as well as pyochelin biosynthesis proteins.[14] The multidrug resistant strain PA14 also showed enhanced transcription of the highly virulent T3SS effectors (*exo*, *pop*, *pcr*) and phenazine biosynthesis at 37 °C.[13] In the pathogen PA254 used in this study, however, certain T3SS effectors (i.e. PemB) and *rhlR*-dependent factors (i.e. WbpX) were significantly higher at 25 °C, which was not observed in model strains (see **Fig 2b**). In addition, host temperatures did not affect phenazine or pyochelin biosynthesis in PA254 (see **Fig 3** and **Dataset S1**).

At ambient temperatures, PAO1 upregulates the virulence agents protease IV,[57] pyoverdine (Pvd),[14] and T1SS and T2SS products (e.g. *apr*).[58] While PA254 also shows highest Pvd and Apr abundances at low temperatures, protease IV expression was not temperature-regulated. Overall, the virulence factors produced PA254 were distinct from PAO1/PA14, and equally distributed between low and high temperatures (see **Fig 2a**). This suggests that PA254 may express more virulence proteins at ambient conditions than these clinical strains, and that *rhlR* quorum sensing control and type 3 secretion are gradually integrated over the temperature spread instead of localized to host conditions. Supporting this, the pathways for quorum sensing were strongly enriched throughout 25-40 °C (see **Fig 3b**).

Similar to virulence, proteins associated with biofilm formation in PA254 partitioned equally between Pel-based initial biofilms at moderate (25-35 °C) and Alg-dominated chronic biofilms at high (37-42 °C) temperatures (see **Fig 2b**). In accord, the demand of sugar precursors for the complex synthesis of alginate biofilms is reflected in the increasing starch and sucrose pathway (see **Fig 3**). Both swarming and twitching, as well as pathway enrichment, confirmed biofilm processes throughout various temperatures. This contrasts with PAO1, PA14, and several clinical isolates, which display the highest biofilm mass and EPS producing genes (*pel*, *psl, alg*) at 20 °C.[16, 59] While biofilms are also formed above that, low temperature results in distinct biofilm architecture and protein and carbohydrate enrichment;[6, 59] indeed, biofilm surfaces in the environment or on skin/wound tissue often experience colder temperatures than internal infections.[17] Out of seven strains tested, only PAO1’s biofilm formation and alginate production partially recovered at 37 °C,[16] hence still below PA254’s equal distribution.

By modulating its proteome, PA254 transitions gradually between a) an ambient-temperature virulent state characterized by a subset of type 1-3 secreted factors, pyoverdine production, exopolysaccharide-rich biofilms, and a strong proportion of planktonic swimmers and surface swarmers; and b) a host-associated virulent state characterized by a subset of highly potent type 2-3 secreted factors, alginate-dominated biofilms, and a strong proportion of surface twitching motility. Besides strain-specific differences, our fine-stepped temperature gradient might provide high enough resolution for this continuous transition to emerge.

### Thermal reprogramming of macro- and micronutrient metabolism

Temperature resulted in the restructuring of major metabolic pathways with pronounced changes surrounding a host-adapted lifestyle. Above 37 °C, PA254 shifts from the degradation of complex environmental xenobiotics and sugar acids to the metabolism of host-derived carbon sources such as glycans, fatty acids, and reduced small molecules (**Fig 3**). A similar trend was observed in other *Pseudomonas* infections through transcriptomics,[60] further confirming host-like conditions of PA254 at these temperatures. Furthermore, changes in the glyoxylate shunt at 35-37 °C are a common signature of pathogen adaptation to host nutrients, oxidative stress, and biofilm conditions.[61]

The negative enrichment of glycine, serine, threonine, histidine and tyrosine metabolism at 37 °C (**Fig 3**) was almost exclusively driven by catabolic and degrading enzymes (*Supplemental Dataset S3*), indicative of amino acid conservation in limited host environments. Supporting this, tyrosine and histidine metabolic gene clusters were found essential for *Pseudomonas* proliferation in surgical wounds,[30] and serine, glycine, and threonine indeed poorly available in the host.[60] These changes are accompanied by a shift from environmental nitrate consumption to reduced nitrite and nitrous oxide typical for oxidative stress and biofilm conditions in the host (see **Fig 2f**).[62]

Uptake of alternative Fe sources was strongly coupled to 37 °C in PA254, which is in agreement with the coupling of Fe-dependent biofilm processes to 37 °C.[34] In addition, previous studies have demonstrated the aerobic metal starvation response of PAO1 and PA14 to differ from the anaerobic response.[35] For example, Fe-acquiring *phuR* was only more abundant under anaerobic metal scarcity,[35] consistent with elevated iron demand under anoxia.[10] Here, significant changes in PhuR abundance together with partial denitrification and biofilm formation indicators suggest existing redox transitions in the PA254 cultures across temperatures, and highlights its elaborate metabolic control within hypoxic gradients commonly encountered in host-associated niches[11] rather than strict oxic or anoxic conditions. *P. aeruginosa*’s ability to rapidly deplete oxygen from even aerated media[63] is consistent with the potential generation of microaerophilic conditions in this study. Still, growth here was likely not completely anaerobic, illustrated by the very low abundance of proteins PA254 highly expresses under true anoxia, such as the class III RNR NrdD, the response regulator CopR, the denitrification protein NorD, and the copper-binding protein NosD (< 10 total spectral counts).[10]

In a transcriptomics study, the *P. aeruginosa* strain PAO1 exhibited slightly different trends to the changes outlined above. A temperature comparison showed less abundant transcripts for the metabolism of starch, sucrose, alcohol, and pyoverdine, and more abundant transcripts for pyochelin metabolism, TCA cycle, and glucose assimilation at 37 °C vs. 22 °C.[14] These differences might stem from the two individual strains used, from the 5-fold less carbon load in our culturing conditions, or from the methodological differences of RNA-versus protein-based ‘omics.[64]

Overall, these trends point toward a host-adapted state of carbon sparing, energy conservation, redox balance control under partial denitrification, and metabolic maintenance under chronic biofilm conditions.

### Decoupling of cobalamin biosynthesis from cellular reservoirs

Across biofilm formation and depth, *P. aeruginosa* transitions from oxygenated to anoxic conditions.[11] Earlier studies have shown that ribonucleotide reductases (RNRs) are vital for anaerobic growth, and that the cobalamin-dependent class II RNR (NrdJ) dominates in biofilms.[65] Under oxygen deficiency, however, several studies confirm downregulation of the O_2_-dependent cobalamin biosynthesis *cob* gene cluster,[30, 56] and NrdJ-dependent *P. aeruginosa* mutants to be growth deficient, presumably due to their lack of genes for O_2_-independent B_12_ production.[29] This led to the hypothesis that aerobic or microaerophilic B_12_ synthesis via the CobN enzyme in upper layers of the biofilm might accumulate enough cobalamin to sustain anoxic biofilm layers through inward diffusion.[31]

Contrary to that assumption, previous metalloproteomics analyses of PA254 revealed the cobalamin biosynthesis protein CobN to be more abundant under oxygen-depleted conditions.[10] Here, CobN was constant throughout the temperature gradient independent of changes in biofilm or redox marker proteins, with a significant decline only under heat stress (**Dataset S1**). Insufficient B_12_ in standard laboratory media (i.e. LB) can limit anaerobic *P. aeruginosa* growth.[29, 56, 66] PA254 was still not at maximum B_12_ production in our 1 μM Co-supplemented growth media (i.e., LB/2), based on >2-fold higher cellular B_12_ with >1 μM Co (**Fig 5a**), but baseline Co perhaps sustained CobN expression. Nonetheless, protein abundance of the overall cobalamin biosynthetic pathway still declined above 22 °C (see **Fig 4f**), suggesting that CobN abundance alone might not be an accurate proxy for B_12_ production. CobN also did not correlate with NrdJ, which was significantly less abundant at 35-42 °C compared to environmental temperatures (**Fig S10**). Lastly, a lower abundance of NrdJ at 35-42 °C with sustained biomass and constant cellular B_12_ could also suggest higher enzymatic efficiencies with increasing temperatures in order to maintain B_12_ biosynthesis and class II RNR activity;[67] a similar decline of NrdAB at these temperatures supports this assumption (**Fig S10**).

Increasing enzymatic activity of B_12_ biosynthesis can also be inferred via kinetic considerations. The change of cellular B_12_ over time (at stationary phase at a specific temperature) is composed of the B_12_-dependent import (B_12-trans_) rate and the Co-dependent biosynthesis (B_12-syn_) rate, minus the rate of B_12_ loss (B_12-loss_) due to efflux, degradation, and other processes (**Equation 1**). From the experiments conducted in this study, we can observe that the change of B_12_ over time approaches zero (**Fig 5b**), with the terms for import *k_1_*[B_12-trans_][B_12_] likely being small (non-detectable B_12_ in fresh medium) and for synthesis *k_2_*[B_12-syn_][Co] synthesis comparatively higher (biosynthesis from added cobalt, **Fig 5a**). The loss term *k_3_*[B_12-loss_] is also considered small or constant, since no release to the media or decline over time was found (**Fig S13**). Therefore, it can be concluded that in case of sufficient [Co] and declining [B_12-syn_], *k_2_* would have to increase with temperature in order to maintain negligible change in (1), as illustrated in **Fig 5c**. Experiments with BtuB-mutants or ^57^Co-labeled B_12_ could be used to further discern these terms and the conceptual model for cellular B_12_ metabolism.

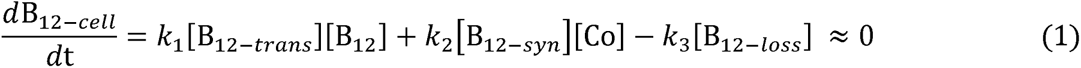

Overall, this dataset shows partitioning into a subset of temperature-independent cobalamin parameters (B_12_ transporters, B_12_-dependent methionine synthase, intracellular B_12_) and a subset of temperature-regulated proteins (B_12_ biosynthesis, B_12_ recycling, B_12_-dependent RNR), with the latter having offset temperature thresholds. We assume that due to a decline of cobalamin biosynthetic proteins, the organism prioritizes certain B_12_ functions (MetH; no change with temperature, **Fig 4f**) at the expense of others (NrdJ; decline with temperature, **Fig 4f**). That MetH abundances in PA254 exceed those of B_12_-independent MetE across temperatures (**Dataset S1**) further confirms that B_12_ remains important for cellular processes throughout temperature- and oxygen-dependent proteome reorganization. Other mechanisms for keeping the cellular cobalamin reservoir constant might be efficient recycling mechanisms, temperature-enhanced enzymatic rates, or storage of B_12_ produced before stationary phase. Since the vitamin B_12_ biosynthesis pathway shows very low abundance at human body temperatures (**Fig 4f**), *P. aeruginosa* is possibly susceptible to Co nutritional immunity strategies by the host, thus creating new avenues for potential drug discovery.

## Conclusion

By combining global proteomics with direct measurements of intracellular cobalamin across seven temperature treatments, this study provides an integrated view of how temperature shapes the functional landscape of *Pseudomonas aeruginosa* PA254. It was observed that virulence, motility, metabolism, and metal homeostasis were responsive to a gradient between environmental and host-relevant temperatures. Based on the high-resolution temperature curve, it is evident that PA254 expresses various proteins involved in biofilm formation and virulent secretion systems throughout, rather than switching between two discrete states. Importantly, the thermal sensitivity of iron acquisition and cobalamin synthesis were decoupled from cellular cobalamin pools, which could contribute to future nutritional immunity research strategies. In a previous transcriptomic study, 6.4% of the *Pseudomonas aeruginosa* genome was found to be differentially regulated between 22 and 37 °C.[14] Between the same treatments, the proteomics approach applied here uncovers 8.9% of differentially abundant proteins among the entire proteome; thus it represents an important complementary tool for systems biology analysis and n promising contributor for future whole cell modeling studies. Follow-up efforts should integrate temperature gradients with the direct manipulation of metal content and control of oxygen availability for this pathogen. Overall, the large scale and coordinated shifts in *P. aeruginosa*’s proteome in response to a host-to-environment temperature gradient reflect this microbe’s extraordinary adaptability and system-level control to varying external conditions.

## Acknowledgements

We thank Fadime Stemmer for assistance with Python coding for visualization.

## Author contributions

V.S.: Data Curation, Formal Analysis, Investigation, Methodology, Visualization, Writing – original draft. M.R.M.: Data Curation, Methodology, Resources. A.S.: Investigation. E.A.W.: Data Curation. M.A.S.: Conceptualization, Funding Acquisition, Project Administration, Resources, Supervision, Writing – review & editing.

## Supporting Information

**Figure S1.** Temperature-dependent growth curves of *Pseudomonas aeruginosa* PA254 replicates cultivated at environmental (22-30 °C), host-associated (35-37 °C), and heat stress conditions (40-42 °C).

**Figure S2.** Features per individual sample (number of unique proteins) for *Pseudomonas aeruginosa* PA254 grown at different temperatures, generated using Fragpipe Analyst.[41]

**Figure S3.** Coefficient of Variation for replicate cultures of *Pseudomonas aeruginosa* PA254 grown at different temperatures, generated using Fragpipe Analyst.[41]

**Figure S4.** Pairwise Pearson correlation for replicate cultures of *Pseudomonas aeruginosa* PA254 grown at different temperatures, generated using Fragpipe Analyst.[41]

**Figure S5.** Log2-centered intensity for replicate cultures of *Pseudomonas aeruginosa* PA254 grown at different temperatures, generated using Fragpipe Analyst.[41]

**Figure S6.** Principle component analysis (PCA) plot for replicate cultures of *Pseudomonas aeruginosa* PA254 grown at different temperatures, generated using Fragpipe Analyst.[41]

**Figure S7.** Counts of differentially expressed proteins (Benjamini-Hochberg adjusted *p* < 0.05) for *Pseudomonas aeruginosa* PA254 grown at different temperatures.

**Figure S8.** Volcano plot of up-(red) and downregulated (blue) proteins (unadjusted *p* < 0.05) for *Pseudomonas aeruginosa* PA254 grown at 42 versus 22 °C.

**Figure S9.** Distribution of differentially expressed proteins (Benjamini-Hochberg adjusted *p*-value < 0.05) within the KEGG pathways of *Pseudomonas aeruginosa* PA254 grown at 22 and 42 °C.

**Figure S10.** Spectral counts of the ribonucleotide reductases NrdA/B (a) and NrdJab, NrdD (b) expressed by for *Pseudomonas aeruginosa* PA254 grown at different temperatures.

**Figure S11.** Molecular structure of the cobalamin or vitamin B12 analogs cyanocobalamin (R = CN), methylcobalamin (R = Me), and hydroxocobalamin (R = OH).

**Figure S12.** Intracellular concentrations of cyanocobalamin (CN-B12), methylcobalamin (Me-B12), and hydroxocobalamin (OH-B12) within *Pseudomonas aeruginosa* PA254 grown at different temperatures.

**Figure S13.** Intracellular concentrations of cyanocobalamin (CN-B12) as proxy during the growth curve of *Pseudomonas aeruginosa* PA254 at 40 °C.

**Table S1.** Abbreviations and descriptions for the temperature-dependent expression of proteins by *Pseudomonas aeruginosa* PA254 depicted in **Figs 2** and **4**.

**Dataset S1.** Differentially expressed proteins and their abundances in *Pseudomonas aeruginosa* PA254 across a temperature gradient from 22-42 °C.

**Dataset S2.** *Pseudomonas aeruginosa* PA254 proteins and their corresponding cellular pathways of the Kyoto Encyclopedia for Genes and Genomes (KEGG) Pathway Database.

**Dataset S3.** Results of *Pseudomonas aeruginosa* PA254 Overrepresentation Analysis (ORA) and Gene Set Enrichment Analysis (GSEA) at different temperature comparisons.

