## Supplementary Material for "Proteomics reveal temperature-coupled cobalamin homeostasis and pathogenicity in *Pseudomonas aeruginosa*"

**For**

**Content	Pages**

**Supplemental Figures S1-S13**	02-14

**Supplemental Table S1**	15-18

**References**	19

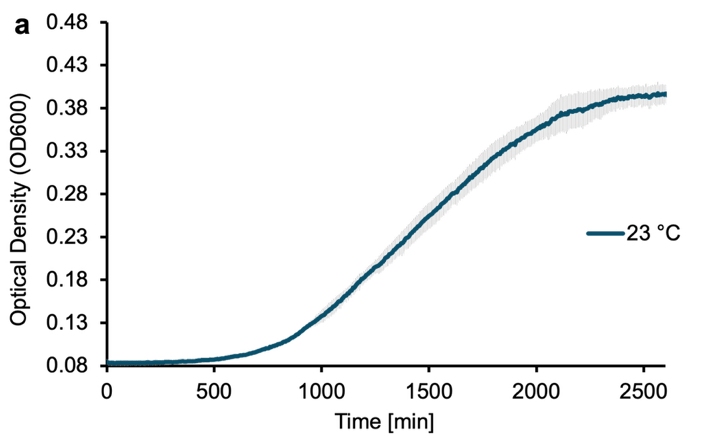

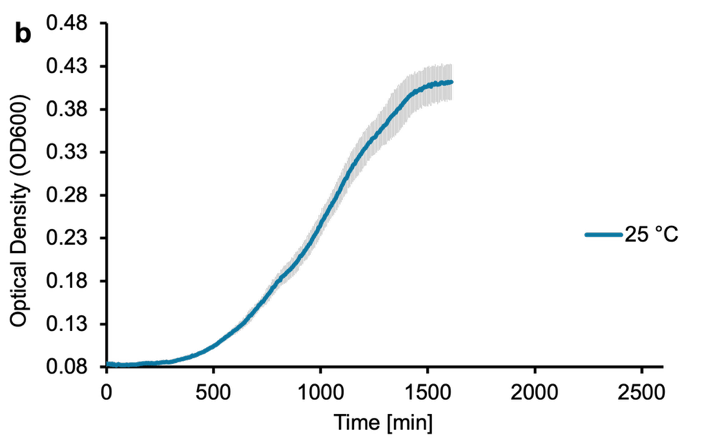

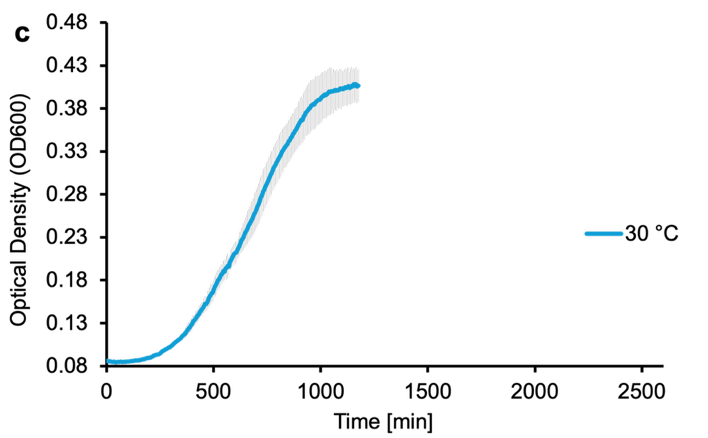

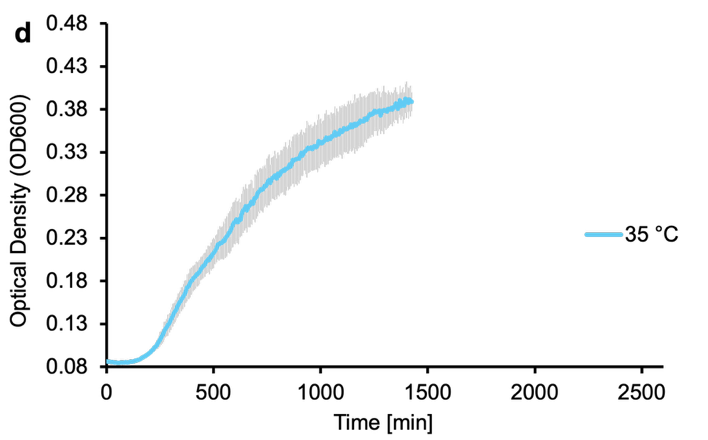

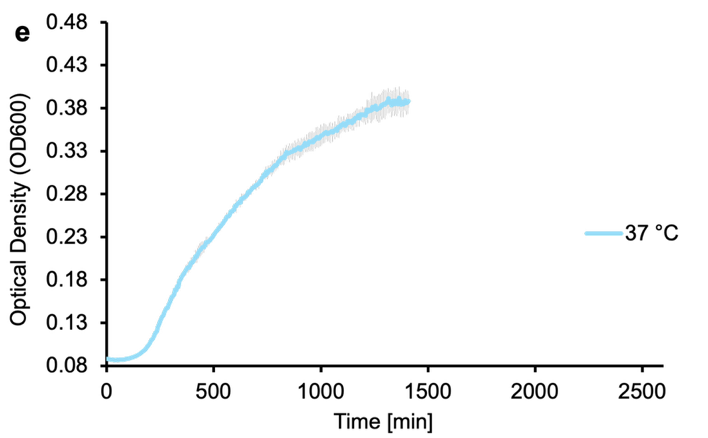

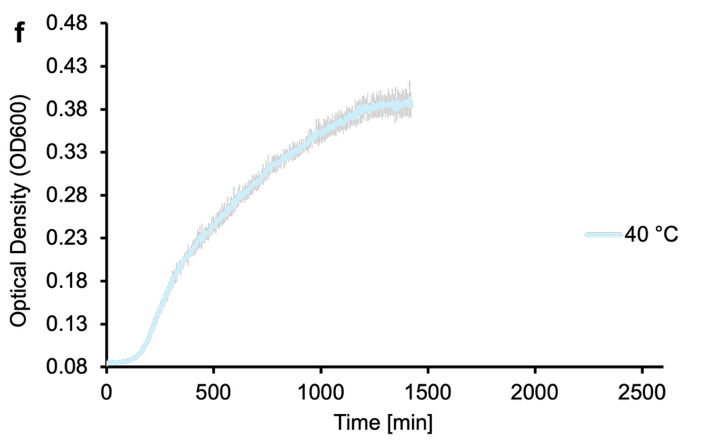

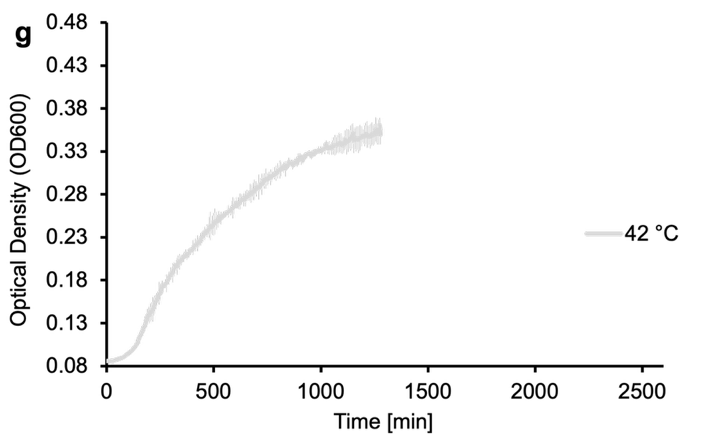

**Figure S1.** Temperature-dependent growth curves of *Pseudomonas aeruginosa* PA254 cultivated at environmental (22-30 °C), host-associated (35-37 °C), and heat stress conditions (40-42 °C), with standard deviation of replicates (*n* = 2 or 3).

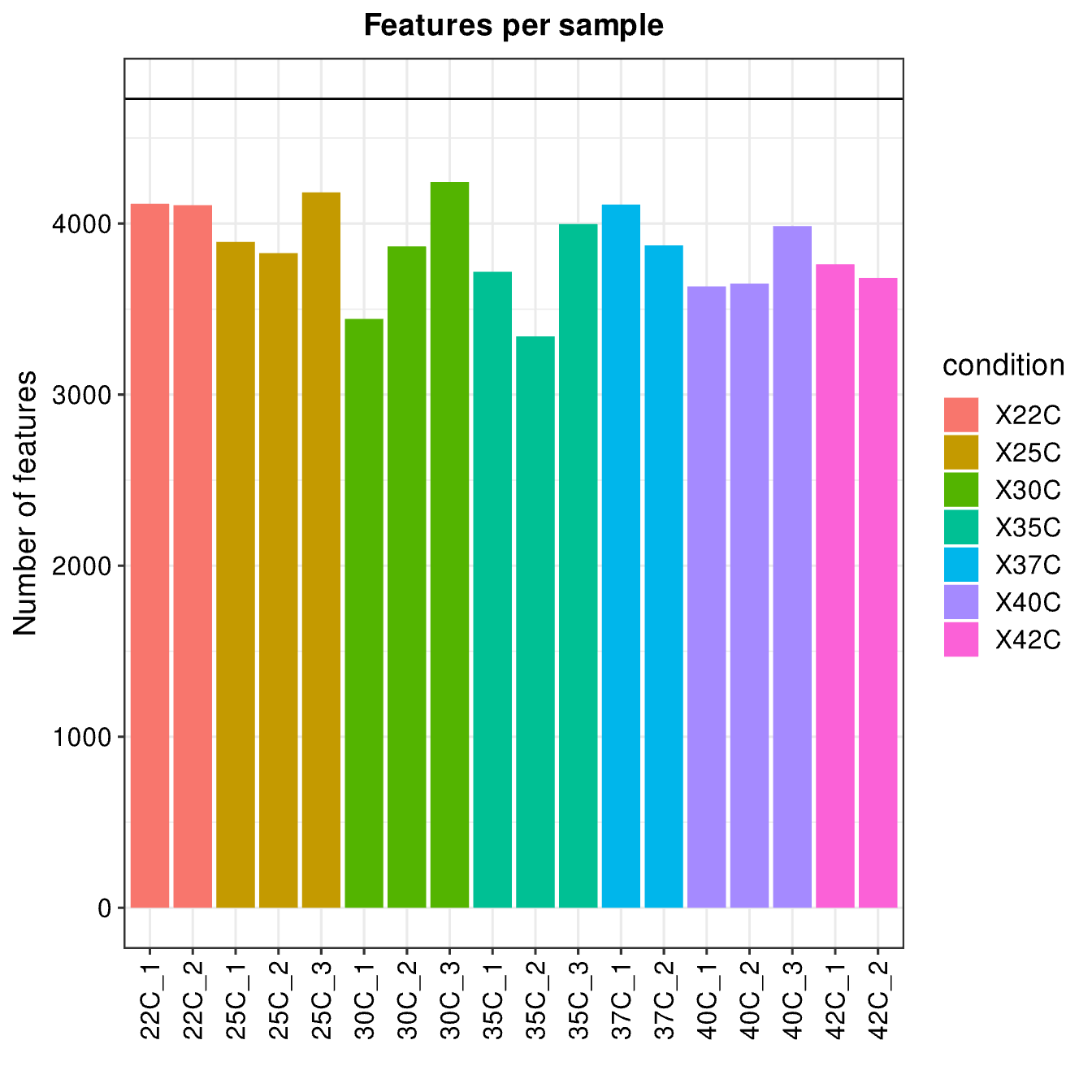

**Figure S2.** Features per individual sample (number of unique proteins) for *Pseudomonas aeruginosa* PA254 grown at different temperatures, generated using Fragpipe Analyst.[1]

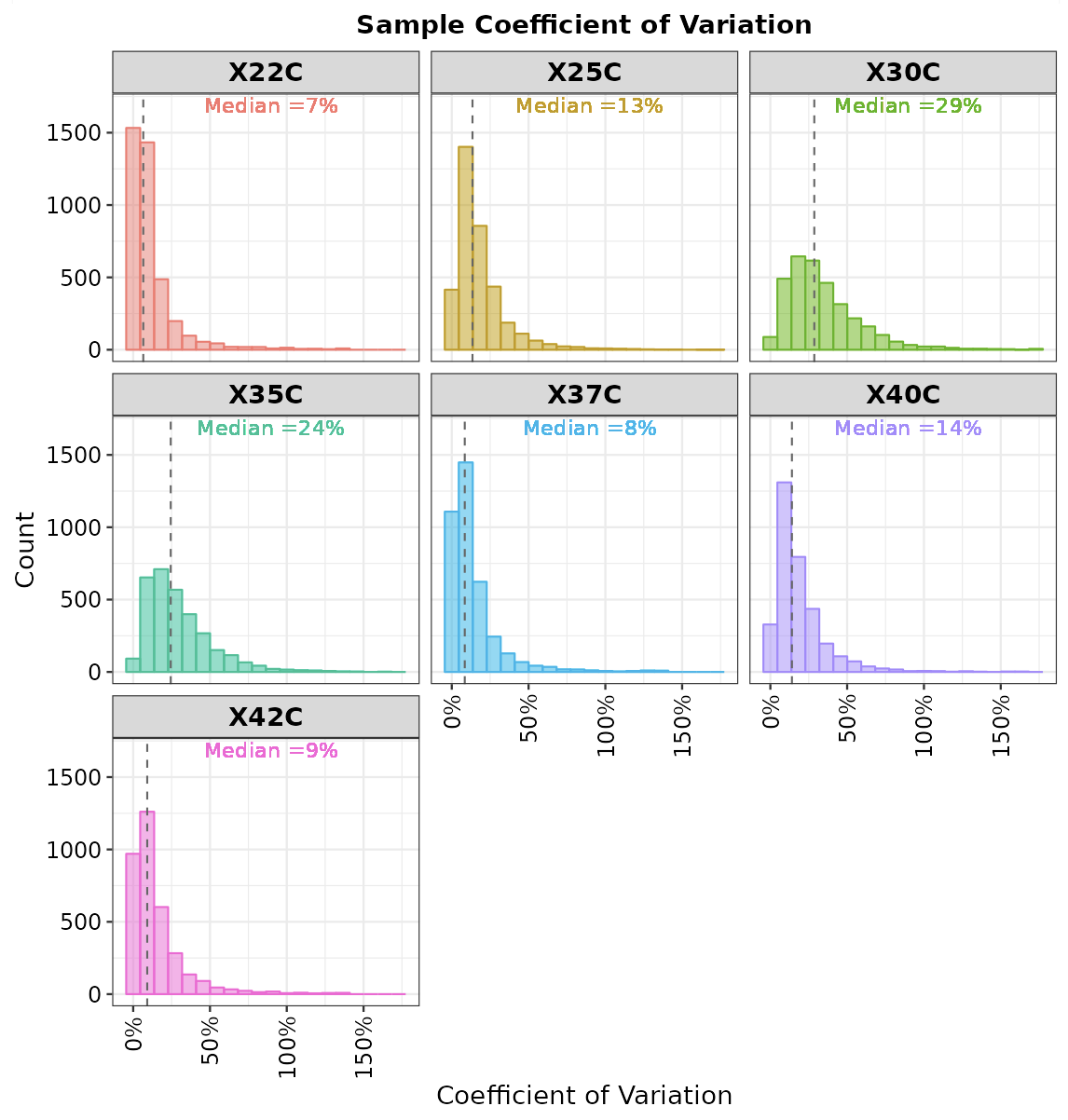

**Figure S3.** Coefficient of Variation for replicate cultures of *Pseudomonas aeruginosa* PA254 grown at different temperatures, generated using Fragpipe Analyst.[1]

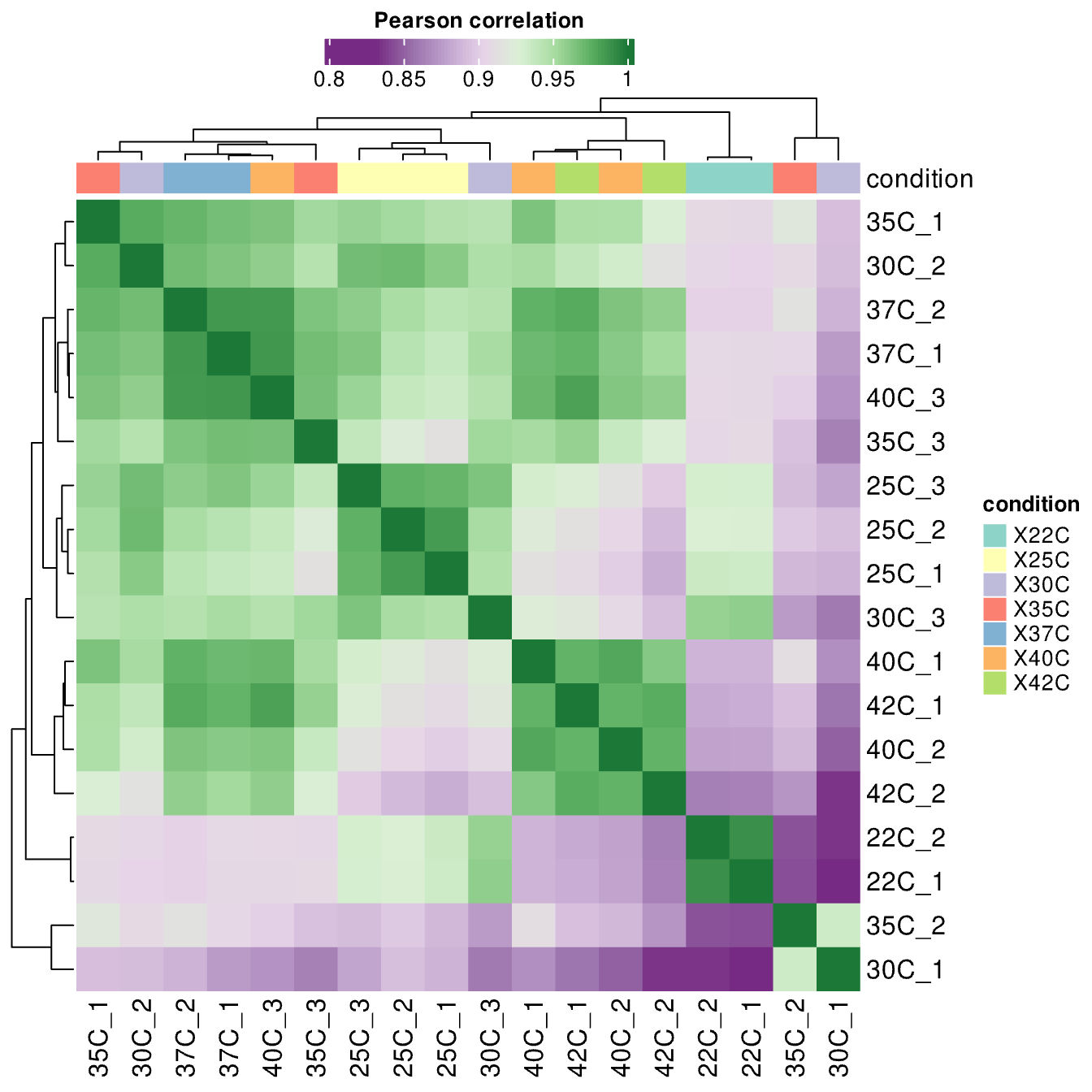
**Figure S4.** Pairwise Pearson correlation for replicate cultures of *Pseudomonas aeruginosa* PA254 grown at different temperatures, generated using Fragpipe Analyst.[1]

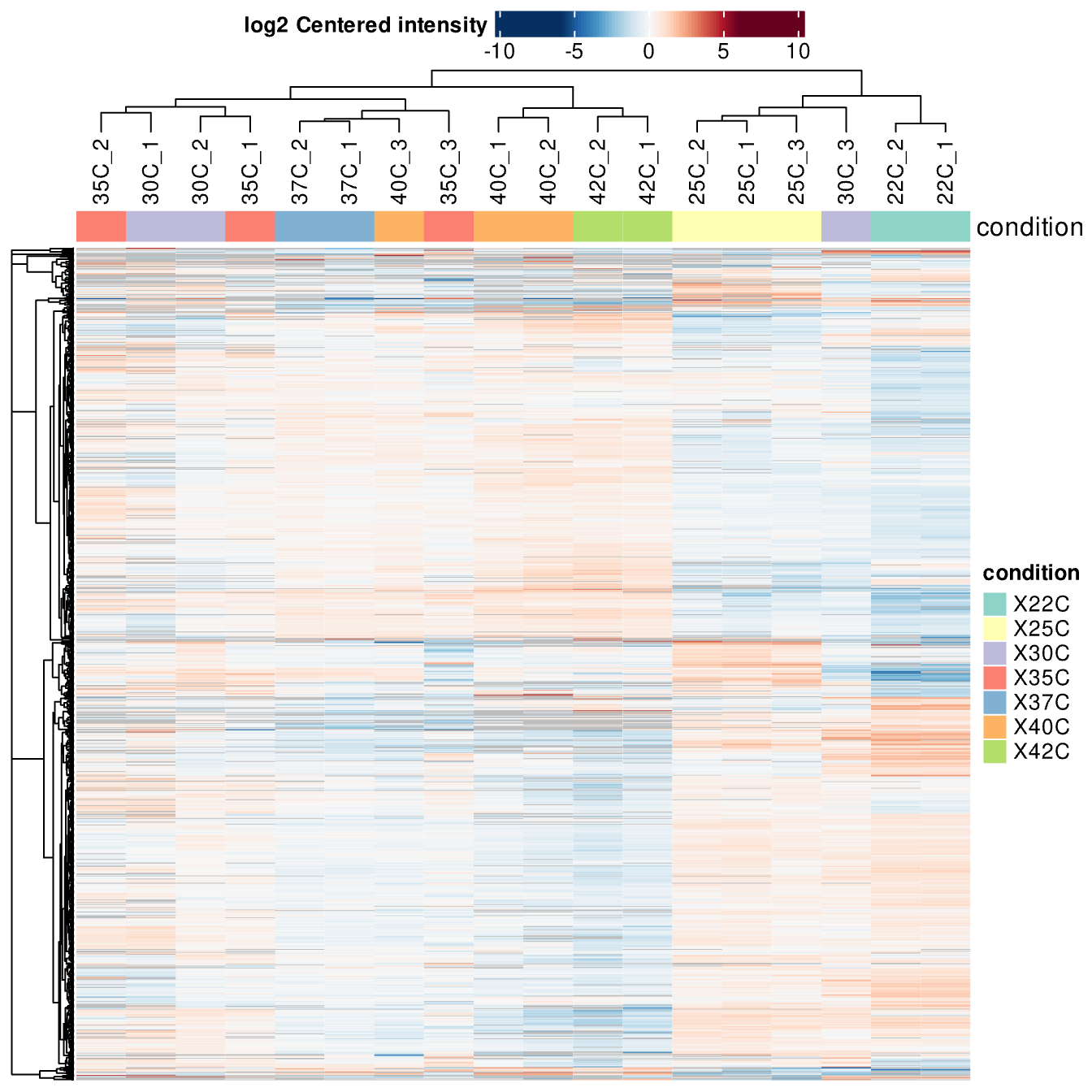
**Figure S5.** Log2-centered intensity for replicate cultures of *Pseudomonas aeruginosa* PA254 grown at different temperatures, generated using Fragpipe Analyst.[1]

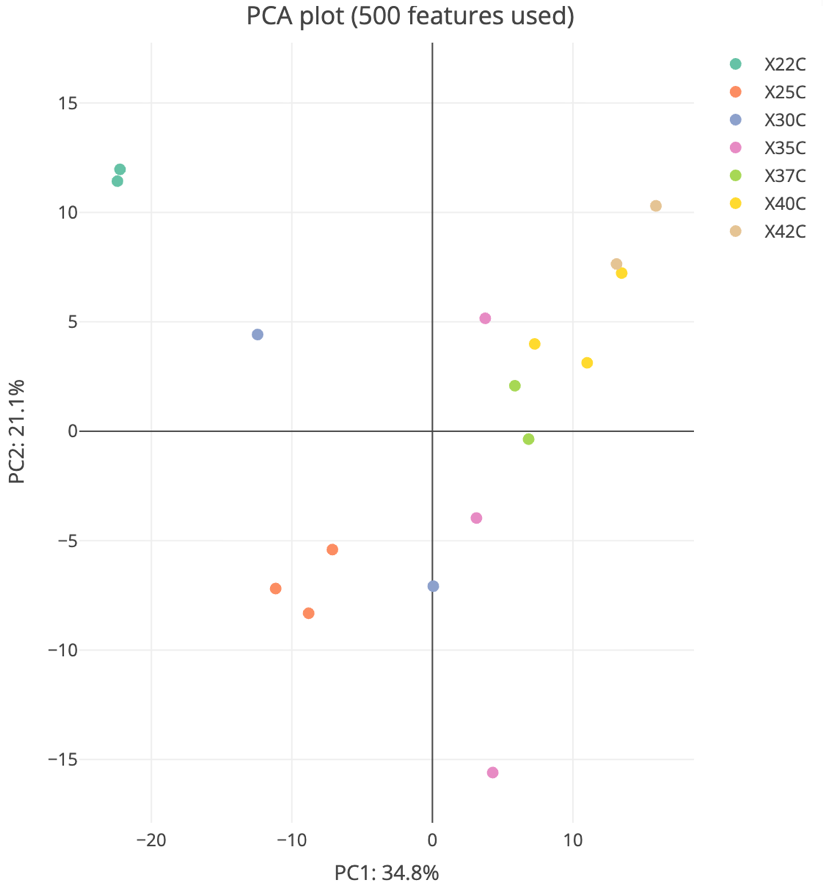

**Figure S6.** Principle component analysis (PCA) plot for replicate cultures of *Pseudomonas aeruginosa* PA254 grown at different temperatures, generated using Fragpipe Analyst.[1]

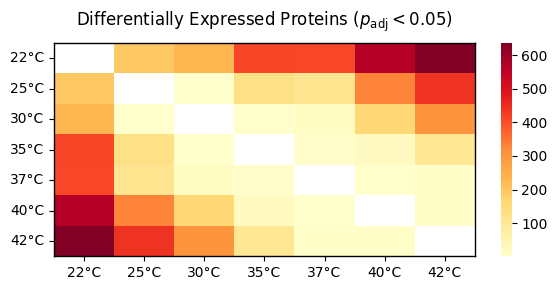

**Figure S7.** Counts of differentially expressed proteins (Benjamini-Hochberg adjusted *p* < 0.05) for *Pseudomonas aeruginosa* PA254 grown at different temperatures.

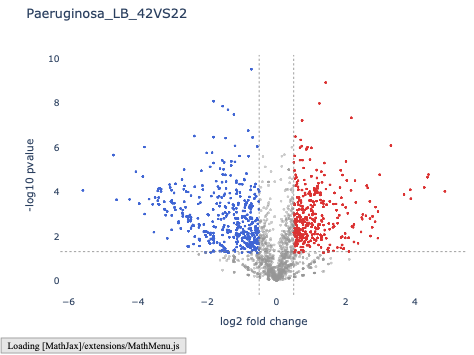

**Figure S8.** Volcano plot of up- (*red*) and downregulated (*blue*) proteins (unadjusted *p* < 0.05) for *Pseudomonas aeruginosa* PA254 grown at 42 versus 22 °C.

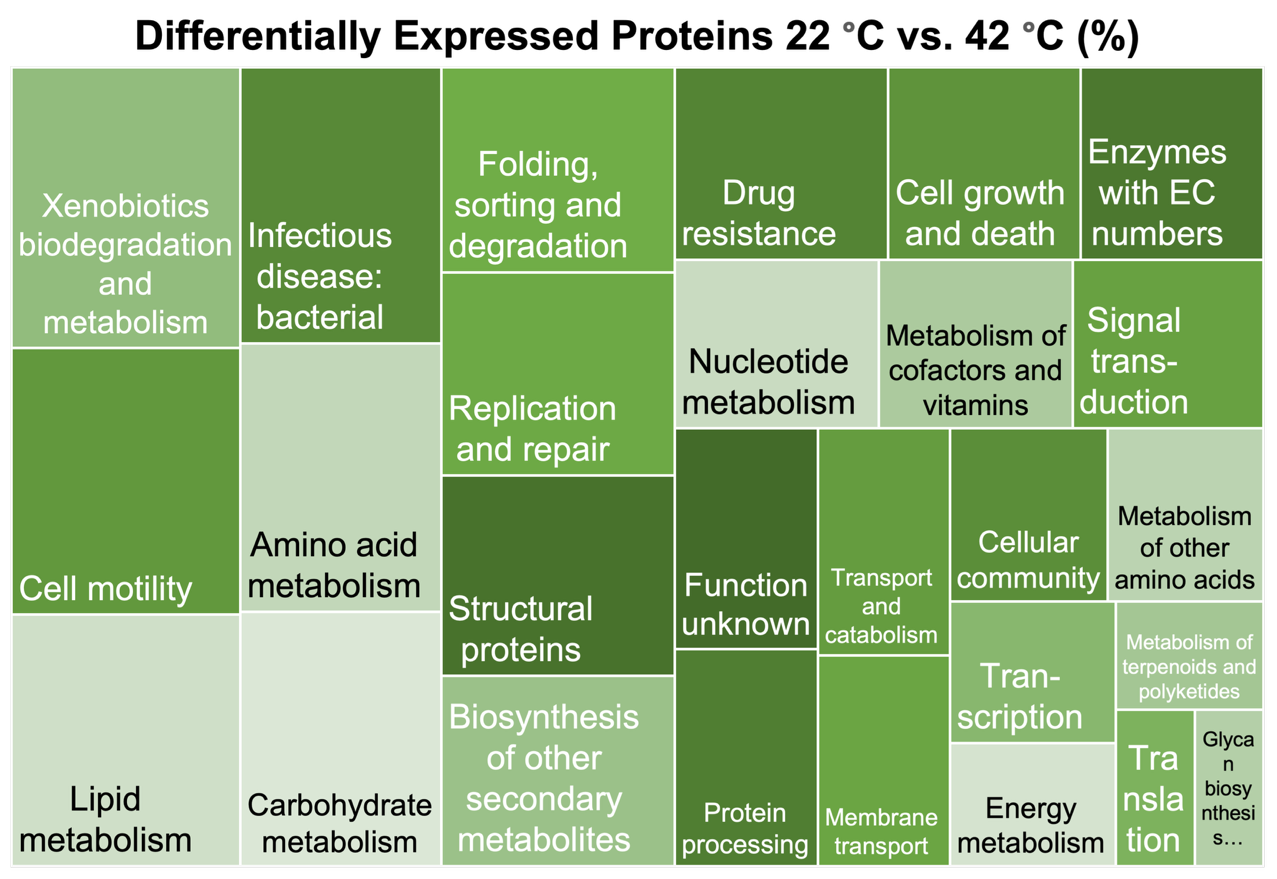

**Figure S9.** Distribution of differentially expressed proteins (Benjamini-Hochberg adjusted *p*-value < 0.05) within the KEGG pathways of *Pseudomonas aeruginosa* PA254 grown at 22 and 42 °C.

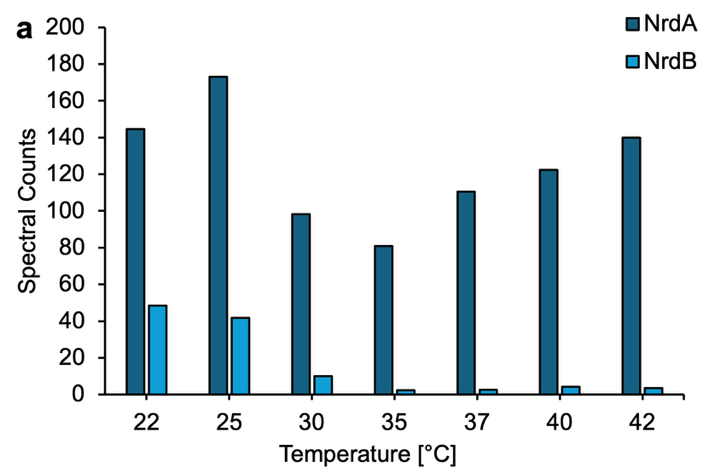

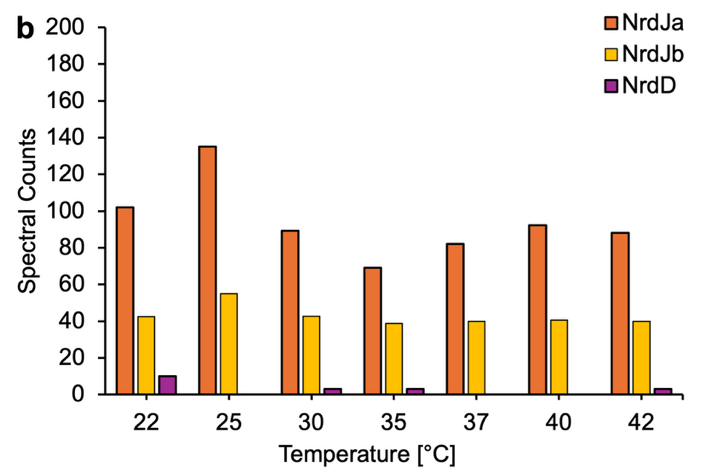

**Figure S10.** Spectral counts of the ribonucleotide reductases NrdA/B **(a)** and NrdJab, NrdD **(b)** expressed by for *Pseudomonas aeruginosa* PA254 grown at different temperatures.

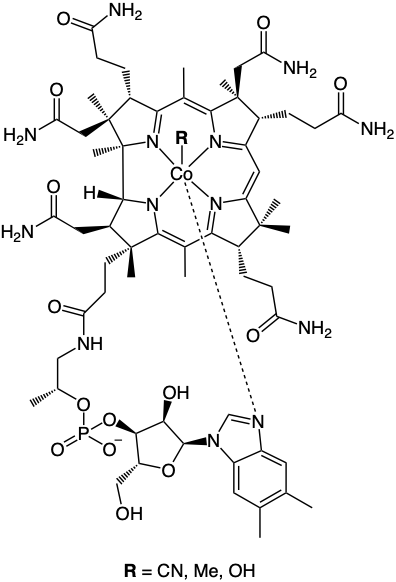

**Figure S11.** Molecular structure of the cobalamin or vitamin B12 analogs cyanocobalamin (R = CN), methylcobalamin (R = Me), and hydroxocobalamin (R = OH).
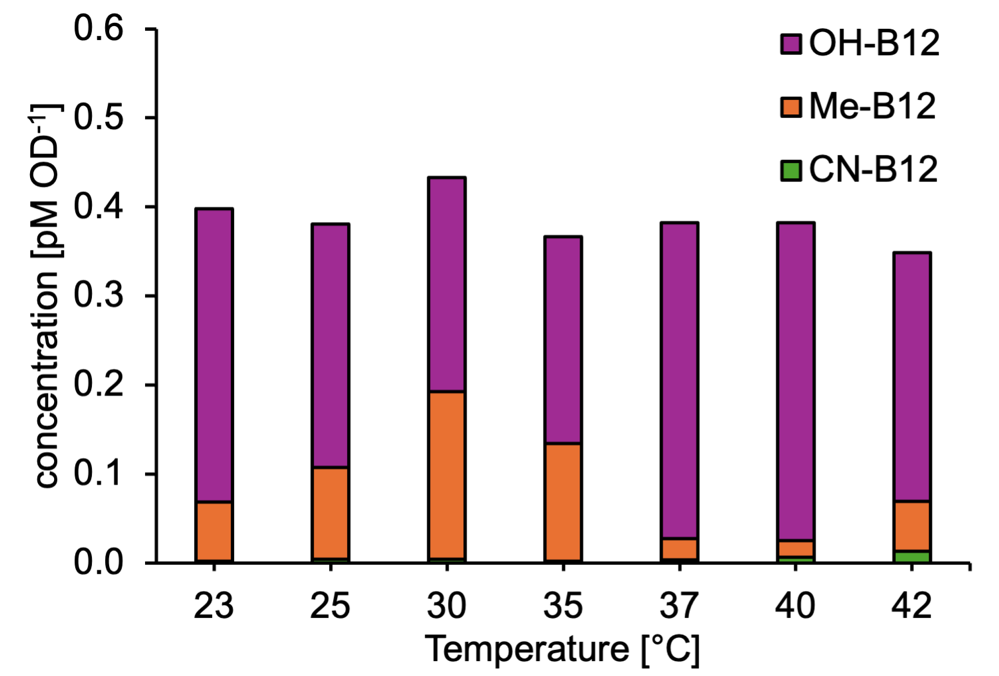

**Figure S12.** Intracellular concentrations of cyanocobalamin (CN-B12), methylcobalamin (Me-B12), and hydroxocobalamin (OH-B12) within *Pseudomonas aeruginosa* PA254 grown at different temperatures.

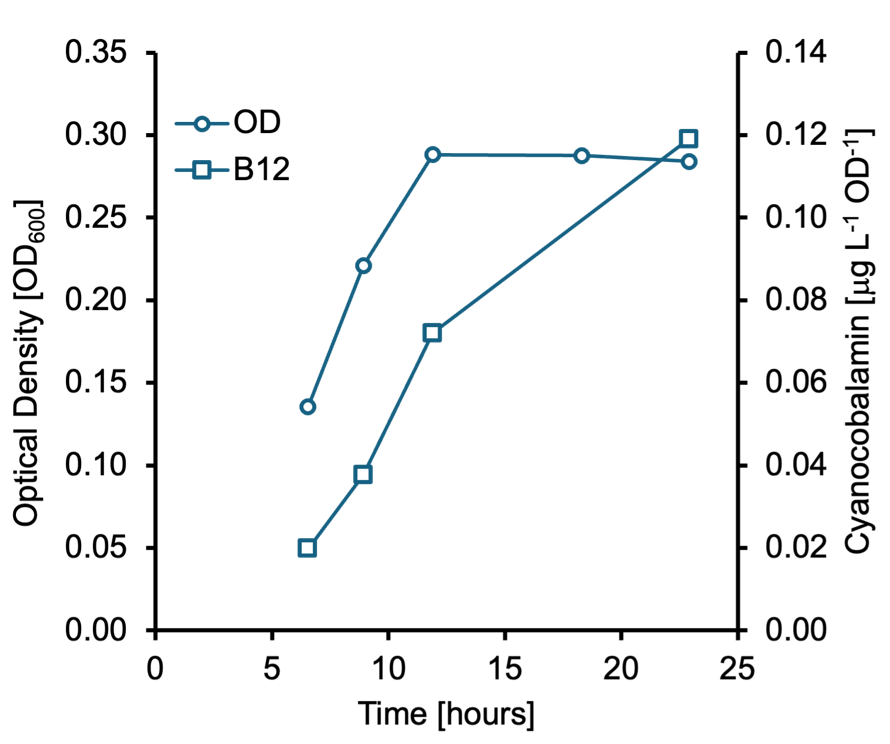

**Figure S13.** Intracellular concentrations of cyanocobalamin (CN-B12) as proxy during the growth curve of *Pseudomonas aeruginosa* PA254 at 40 °C.

**Table S1.** Abbreviations and descriptions for the temperature-dependent expression of proteins by *Pseudomonas aeruginosa* PA254 depicted in **Figs 2** and **4**.

| **Protein** | **Category** | **Description (KEGG Orthology)** |
| --- | --- | --- |
| *Heat Shock* | | |
| ClpB | Chaperone | K03695 Clp protease ATP-binding subunit ClpB |
| GroEL | Chaperone | K04077 chaperonin GroEL |
| DnaJ | Chaperone | K09522 DnaJ heat shock protein family (Hsp40) member |
| DnaK | Chaperone | K04043 molecular chaperone DnaK |
| *Virulence Factors* | | |
| LipA | Exotoxin | K01046 lipA; triacylglycerol lipase |
| PemB | Exotoxin | no KO assigned \| PemB, type III secretion effector |
| AprE | Surface Structure | K01406 alkaline metalloproteinase; serralysin |
| AprF | Surface Structure | no KO assigned \| Type I Secretion System |
| WbpX | Surface Structure | K24655 wbpX; glycosyltransferase WbpX |
| PvdF | Secreted Factor | no KO assigned \| pvdF; pyoverdine synthetase F |
| PvdJ | Secreted Factor | no KO assigned \| pvdJ; pyoverdine biosynthesis protein PvdJ |
| PvdL | Secreted Factor | no KO assigned \| pvdL; peptide synthase |
| PvdQ | Secreted Factor | K07116 pvdQ; acyl-homoserine-lactone acylase |
| PvdR | Secreted Factor | K13888 pvdR; pyoverdine biosynthesis protein PvdR |
| PvdA | Secreted Factor | K10531 pvdA; L-ornithine N5-monooxygenase |
| PvdD | Secreted Factor | no KO assigned \| pvdD; pyoverdine synthetase D |
| PvdE | Secreted Factor | K06160 pvdE; pyoverdine biosynthesis protein PvdE |
| PvdT | Secreted Factor | K05685 (RefSeq) pvdT; pyoverdine biosynthesis protein PvdT |
| PvdN | Secreted Factor | no KO assigned \| pvdN; pyoverdine biosynthesis protein PvdN |
| PvdP | Secreted Factor | no KO assigned \| pvdP; pyoverdine biosynthesis protein PvdP |
| LasA | Secreted Factor | K08642 LasA protease |
| LasR | Regulator | K18304 quorum-sensing system regulator LasR |
| ExoU | Exotoxin | K16638 exoenzyme U |
| PopB | Surface Structure | K23475 transloator protein popB |
| PopD | Surface Structure | K23476 translocator outer membrane protein PopD |
| PcrV | Surface Structure | no KO assigned \| type III secretion protein PcrV |
| *Biofilm Formation* | | |
| PslB | EPS matrix | K16011 polysaccharide biosynthesis protein PslB |
| PslC | EPS matrix | K25205 polysaccharide biosynthesis protein PslC |
| PslD | EPS matrix | K20987 polysaccharide biosynthesis/export protein PslD |
| PslG | EPS matrix | K21000 polysaccharide biosynthesis protein PslG |
| MucA | EPS matrix | K03597 sigma factor algU negative regulatory protein MucA |
| MucB | EPS matrix | K03598 sigma factor AlgU regulator MucB |
| AlgB | EPS matrix | K11384 two-component response regulator AlgB |
| AlgR | EPS matrix | K08083 two-component system, response regulator AlgR |
| *Swimming Motility* | | |
| FliC | Flagellum | K02406 fliC; flagellin |
| FliF | Flagellum | K02409 fliF; flagellar M-ring protein |
| **Protein** | **Category** | **Description (KEGG Orthology)** |
| FliG | Flagellum | K02410 fliG; flagellar motor switch protein |
| FliO | Flagellum | K02418 flagellar assembly protein FliO/FliZ |
| FliS | Flagellum | K02422 fliS; flagellar secretion chaperone |
| FlgA | Flagellum | K02386 flgA; flagellar basal-body P-ring formation protein |
| FlgD | Flagellum | K02389 flgD; flagellar basal-body rod modification protein |
| FlgE | Flagellum | K02390 flgE; flagellar hook protein |
| FlgH | Flagellum | K02393 flagellar L-ring protein FlgH |
| FlgI | Flagellum | K02394 flagellar P-ring protein FlgI |
| FlgK | Flagellum | K02396 flagellar hook-filament junction protein FlgK |
| *Twitching Motility* | | |
| PilB | Pilus | K02652 pilB; type IV pilus assembly protein PilB |
| PilG | Pilus | K02657 two-component system response regulator PilG |
| PilH | Pilus | K02658 two-component system response regulator PilH |
| PilM | Pilus | K02662 pilM; type IV pilus assembly protein PilM |
| PilN | Pilus | K02663 pilN; type IV pilus assembly protein PilN |
| PilO | Pilus | K02664 pilO; type IV pilus assembly protein PilO |
| PilQ | Pilus | K02666 pilQ; type IV pilus assembly protein PilQ |
| PilR | Pilus | K02667 two-component system response regulator PilR |
| PilY1 | Pilus | K02674 pilY1; type IV pilus assembly protein PilY1 |
| PilY2 | Pilus | K02675 pilY1; type IV pilus assembly protein PilY2 |
| FimUT | Pilus | K08084 type IV fimbrial biogenesis protein FimT |
| *Denitrification* | | |
| NarK1 | Transporter | K02575 MFS transporter, NNP family, nitrate/nitrite transporter |
| NarK2 | Transporter | K02575 MFS transporter, NNP family, nitrate/nitrite transporter |
| Fhp | Oxygenase | K05916 fhp; nitric oxide dioxygenase |
| ANR | Regulator | K01420 transcriptional regulator, anaerobic regulatory protein |
| DNR | Regulator | K21563 dissimilatory nitrate respiration transcriptional regulator |
| NarG | Reductase | K00370 narG; nitrate reductase/nitrite oxidoreductase, alpha |
| NarH | Reductase | K00371 narH; nitrate reductase/nitrite oxidoreductase, beta |
| NirS | Reductase | K15864 nitrite reductase (NO-forming)/hydroxylamine reductas |
| NorB | Reductase | K04561 norB; nitric oxide reductase subunit B |
| NorC | Reductase | K02305 norC; nitric oxide reductase subunit C |
| YfdC | Transporter | K21990 YfdC formate-nitrite transporter family protein |
| *Iron Uptake* | | |
| FeoC | Transporter | K07490 ferrous iron transport protein C |
| FecA | Transporter | K16091 Fe(3+) dicitrate transport protein |
| FpvA | Transporter | K16088 fpvA; outer membrane ferripyoverdine receptor |
| FpvB | Transporter | K16088 fpvB; second ferric pyoverdine receptor FpvB |
| Fur | Regulator | K03711 fur; ferric uptake regulation protein |
| PhuR | Transporter | K16087 phuR; heme/hemoglobin uptake outer membrane receptor |
| HxuA | Transporter | K16087 hemoglobin/transferrin/lactoferrin receptor protein |
| **Protein** | **Category** | **Description (KEGG Orthology)** |
| PfeA | Transporter | K19611 ferric enterobactin receptor |
| PirA1 | Transporter | K19611 pirA; outer membrane receptor FepA, partial |
| PirA2 | Transporter | K19611 pirA; outer membrane receptor FepA, partial |
| FemA | Transporter | K02014 femA; outer membrane ferric-mycobactin receptor FemA |
| FoxA | Transporter | K02014 foxA; outer membrane ferrioxamine receptor FoxA |
| FiuA | Transporter | K02014 fiuA; outer membrane ferrichrome receptor FiuA |
| *Cobalamin Biosynthesis* | | |
| CobD | Enzyme | K02227 adenosylcobinamide-phosphate synthase CobD |
| CobH | Enzyme | K06042 cobH; precorrin-8X methylmutase |
| CobI | Enzyme | K03394 cobI; precorrin-2 C20-methyltransferase |
| CobJ | Enzyme | K13541 cobalt-precorrin 5A hydrolase / cobalt-factor III methyltransferase / precorrin-3B C17-methyltransferase |
| CobR | Enzyme | K13786 cobR; cob(II)yrinic acid a,c-diamide reductase |
| CobW | Enzyme | K02234 cobalamin biosynthesis protein CobW |
| CysG | Enzyme | K02302 cysG; uroporphyrin-III C-methyltransferase / precorrin-2 dehydrogenase |
| PduO | Enzyme | K00798 cob(I)alamin adenosyltransferase |
| *Cobalamin Usage* | | |
| MetE | Enzyme | K00549 metE; B12-independent methionine synthase |
| MetH | Enzyme | K00548 metH; B12-dependent methionine synthase |
| MetR | Enzyme | K03576 LysR family transcriptional regulator for metE and metH |
| NrdA | Enzyme | K00525 nrdA; ribonucleotide-diphosphate reductase subunit alpha |
| NrdB | Enzyme | K00526 nrdB; ribonucleotide-diphosphate reductase subunit beta |
| NrdJa | Enzyme | K00525 nrdJa; ribonucleoside-diphosphate reductase |
| NrdJb | Enzyme | no KO assigned \| nrdJb; ribonucleoside-diphosphate reductase |
| NrdD | Enzyme | K21636 nrdD; anaerobic ribonucleoside triphosphate reductase |
| *Cobalamin Uptake* | | |
| BtuB1 | Transporter | K16092 vitamin B12 transporter \| tonB-dependent receptor |
| BtuR | Transporter | K19221 cobO; cob(I)yrinic acid a,c-diamide adenosyltransferase |
| BtuB2 | Transporter | K02014 tonB-dependent receptor (iron.B12.siderophore.hemin) |
| BtuF | Transporter | K02016 substrate-binding protein (iron.B12.siderophore.hemin) |
| BtuB3 | Transporter | K02014 tonB-dependent receptor (iron.B12.siderophore.hemin) |
| BtuN | Transporter | no KO assigned \| hypothetical component of the B12 transporter BtuN |
| *Zinc Uptake* | | |
| Zur | Transporter | K09823 Fur family transcriptional regulator, zinc uptake regulator |
| ZnuA | Transporter | K09815 zinc transport system substrate-binding protein |
| ZnuC | Transporter | K09817 zinc transport system ATP-binding protein |
| ZnuD | Transporter | K02014 zinc transport outermembrane recepter protein |
| HmtA | Transporter | no KO assigned \| cation-transporting P-type ATPase |
| PA4066 | Transporter | K09950 uncharacterized protein, putative Zn transporter |
| PA4065 | Transporter | K02004 putative ABC transport system permease protein |
| PA4063 | Transporter | no KO assigned \| putative Zn transporter protein |
| **Protein** | **Category** | **Description (KEGG Orthology)** |
| *Manganese Uptake* | | |
| MntH | Transporter | K03322 mntH; manganese transport protein mntH |
| MntA1 | Transporter | K09815 ABC transport system substrate-binding protein |
| MntC1 | Transporter | K09817 ABC transport system ATP-binding protein |
| MntB1 | Transporter | K09816 ABC transport system permease protein |
| MntA2 | Transporter | no KO assigned \| hypothetical protein |
| *Copper Uptake* | | |
| NosD | Transporter | K07218 nosD; copper-binding periplasmic protein |
| NosY | Transporter | K19341 nosY; Cu-processing system permease protein |
| OprC | Transporter | K02014 oprC; copper transport outer membrane porin OprC |
| SenC | Transporter | K07152 senC; cytochrome c oxidase assembly protein SenC |
| OpdT | Transporter | no KO assigned \| opdT; tyrosine porin OpdT |
| CuPATP1 | Transporter | no KO assigned \| cation-transporting P-type ATPase |

**References.**

1. Hsiao Y *et al.* Analysis and Visualization of Quantitative Proteomics Data Using FragPipe-Analyst. *J Proteome Res*. 2024;23(10):4303–15. doi:10.1021/acs.jproteome.4c00294.
